# Telomerase deficiency in humans is associated with systemic age-related changes in energy metabolism

**DOI:** 10.1101/2022.02.28.481982

**Authors:** Emma Naomi James, Virag Sagi-Kiss, Mark Bennett, Maria Mycielska, Karen-Ng Lee Peng, Terry Roberts, Sheila Matta, Inderjeet Dokal, Jacob Guy Bundy, Eric Kenneth Parkinson

## Abstract

Underlying mechanisms of plasma metabolite signatures of human ageing and age-related diseases are not clear but telomere attrition and dysfunction are central to both. Dyskeratosis Congenita (DC) is associated with mutations in the telomerase enzyme complex (*TERT*, *TERC*, *DKC1*) and progressive telomere attrition. We show extracellular citrate is repressed by canonical telomerase function *in vitro* and associated with DC leukocyte telomere attrition *in vivo;* leading to the hypothesis that altered citrate metabolism detects telomere dysfunction. However, citrate and senescence factors only weakly distinguished DC patients from controls, whereas other tricarboxylic acid cycle metabolites, lactate and especially pyruvate distinguished them with high significance, consistent with further metabolism of citrate and lactate in the liver and kidneys. Citrate uptake in certain organs modulates age-related disease in mice and our data has similarities with age-related disease signatures in humans. Our results have implications for the early diagnosis of telomere dysfunction and anti-senescence therapeutics.

**Highlights:**

- Extracellular citrate is regulated by telomere function *in vitro and in vivo*.
- Dyskeratosis Congenita (DC) is a human disease characterized by systemic telomere attrition, which showed an age-related plasma energetic profile, distinct from age-related disease and that of centenarians.
- The DC profile strikingly out-performed senescence factors in discriminating DC from controls, and pyruvate associated with a low lactate:pyruvate ratio is potentially a useful and cheap minimally invasive diagnostic aid for DC and telomere dysfunction.
- Mechanistically DC systemic metabolism is indicative of a shift to reduced pyruvate dehydrogenase activity, glycolysis and/or increased citrate and lactate production followed by further metabolism in the kidneys and liver.

## Introduction

Cellular senescence is a dynamic process induced by a variety of cellular stresses, including irreparable DNA double strand breaks (IrrDSBs) which accumulate at telomeres following telomerase deficiency and either telomere attrition (d’Adda di Fagagna et al., 2003) or in non-dividing cells due to inadequate DNA repair (Fumagalli et al., 2012; Hewitt et al., 2012). IrrDSBs are resolved by telomerase in dividing cells (Patel et al., 2016; Suram et al., 2012) but not so readily in non-dividing cells.

Telomeres do shorten in a variety of telomerase competent human cells and tissues with chronological age, including squamous epithelia (Nakamura et al., 2002) and leukocytes (Hastie et al., 1990). Additionally, age-adjusted leukocyte telomere length is associated with poor health, especially cardiovascular disease (Epel et al., 2004; Wang et al., 2018), and is reversible upon positive changes in lifestyle (Sellami et al., 2021). Furthermore, the telomerase activator TA-65 has been reported to improve macular degeneration in humans (Dow and Harley, 2016).

In mice, telomere dysfunction induced by complete *Tert* or *Terc* deletion results in impaired mitochondrial biogenesis and function (Sahin et al., 2011), decreased gluconeogenesis, cardiomyopathy, and increased production of reactive oxygen species (ROS). These phenotypes are countered by increased telomerase function (Jaskelioff et al., 2011; Sahin et al., 2011; Tomas-Loba et al., 2008) and excessively long telomeres (Munoz-Lorente et al., 2019), to benefit lifespan and healthspan (Munoz-Lorente et al., 2019; Tomas-Loba et al., 2008). However, the amount of telomere attrition in dividing human cells with age is not as dramatic as that seen in the telomerase *tert*-/- or *terc*-/- mice.

Telomere dysfunction culminates in cellular senescence, which is accompanied by the accumulation of an array of proteins and metabolites outside the cells, known as the senescence-associated secretory phenotype (SASP (Coppe et al., 2008)) and the extracellular senescence metabolome (ESM (James et al., 2015)), respectively. Some metabolites of the ESM are associated with chronological ageing in humans (Auro et al., 2014; Menni et al., 2013), but the underlying mechanisms are largely unknown.

Citrate is particularly interesting because it has been implicated in ageing (Menni et al., 2013; Rogina, 2017), caloric restiction (Rogina, 2017), type 2 diabetes (Birkenfeld et al., 2011; Brachs et al., 2016), blood pressure (Willmes et al., 2021), heart rate (Willmes et al., 2021), memory (Fan et al., 2021) and cancer (Mycielska et al., 2018). Dietary citrate also induces adipocyte inflammation and pre-insulin resistance in mice (Leandro et al., 2016). Apart from its role supplying the TCA cycle, citrate has roles in fatty acid synthesis (Mycielska et al., 2015) and epigenetic regulation via ATP-citrate lyase, acetyl CoA and histone acetylation (Wellen et al., 2009).

Several anti-senescence approaches are now being considered to ameliorate age-related diseases, including senolytic drugs (Campisi et al., 2019), which have shown some promise in clinical trials. However, the non-invasive detection of senescent cells and telomere attrition in human disease is challenging owing to their small number. SASP proteins are not generally detectable, their detectability varies in different disease states (Hickson et al., 2019; Justice et al., 2019; Suvakov et al., 2021; Yousefzadeh et al., 2018) and it is acknowledged that additional plasma biomarkers are needed (reviewed in (Campisi et al., 2019)).

To examine the effect of telomere dysfunction on human metabolites related to cellular senescence or age-related diseases we took advantage of Dyskeratosis Congenita (DC) patient plasma samples. DC patients possess heterozygous (*TERT*/*TERC*) or hemizygous (*DKC1*) rather than homozygous loss of function mutations, and is phenotypically closer to the *terc*+/- mouse than the *tert*-/- or *terc*-/- mouse (Hao et al., 2005).

DC patients display short telomeres in their tissues *in vivo* (Gadalla et al., 2010; Vulliamy et al., 2001) and premature senescence *in vitro* (Mitchell et al., 1999; Sun et al., 2020). DC patients show several markers of aplastic anaemia, skin and nail abnormalities, fibrosis of the lungs and liver, an increased risk of cancer, osteoporosis and more rarely other abnormalities such as cerebellar atrophy (Dokal, 2011). DC patients are a useful study group for the investigation of the effects of telomere dysfunction and senescence on human metabolism because the mechanism of premature senescence is characterised and DC patients do not usually have confounding conditions such as type 2 diabetes (T2D) at an early age (Dokal, 2011; Guo et al., 2011), which would affect metabolism.

We show here that humans with defective telomere maintenance display changes in energy metabolism in the absence of a high senescent cell load and mitochondrial oxidative damage. We also identify metabolites that distinguish DC from controls with high specificity and selectivity that may have diagnostic significance.

## Results

### Citrate is regulated by telomere function and the canonical function of telomerase, independently from inflammatory cytokines *in vitro*

Extracellular citrate (EC) and interleukin 6 (IL-6) both increased in senescent fibroblasts. The catalytic subunit of telomerase, *TERT*, reduced levels of EC (Figure 1A, 1B) in parallel with the frequency of senescence-associated beta galactosidase (Figure 1C, 1D) in both BJ fibroblasts and NHOF-1 cells, but the empty vector and *TERT-HA* (extrachromosomal telomerase functions only) did not. Only the *TERT* transgene was able to increase telomere length (Figures 1E, 1F) despite the fact that *TERT-HA* was expressed (Figure 1G, 1H) and induced telomerase activity in BJ cells (Figure 1I). These data demonstrate that EC is regulated by the canonical function of telomerase and links telomere function to EC accumulation in senescent fibroblasts. Neither *TERT* nor *TERT-HA* reduced IL-6 levels in BJ cells (Figure 1J) under the *in vitro* conditions described here indicating that EC is regulated independently from the inflammatory cytokines. These *in vitro* results gave rise to the hypothesis that plasma citrate is upregulated *in vivo* by telomere attrition.

**Figure 1.**
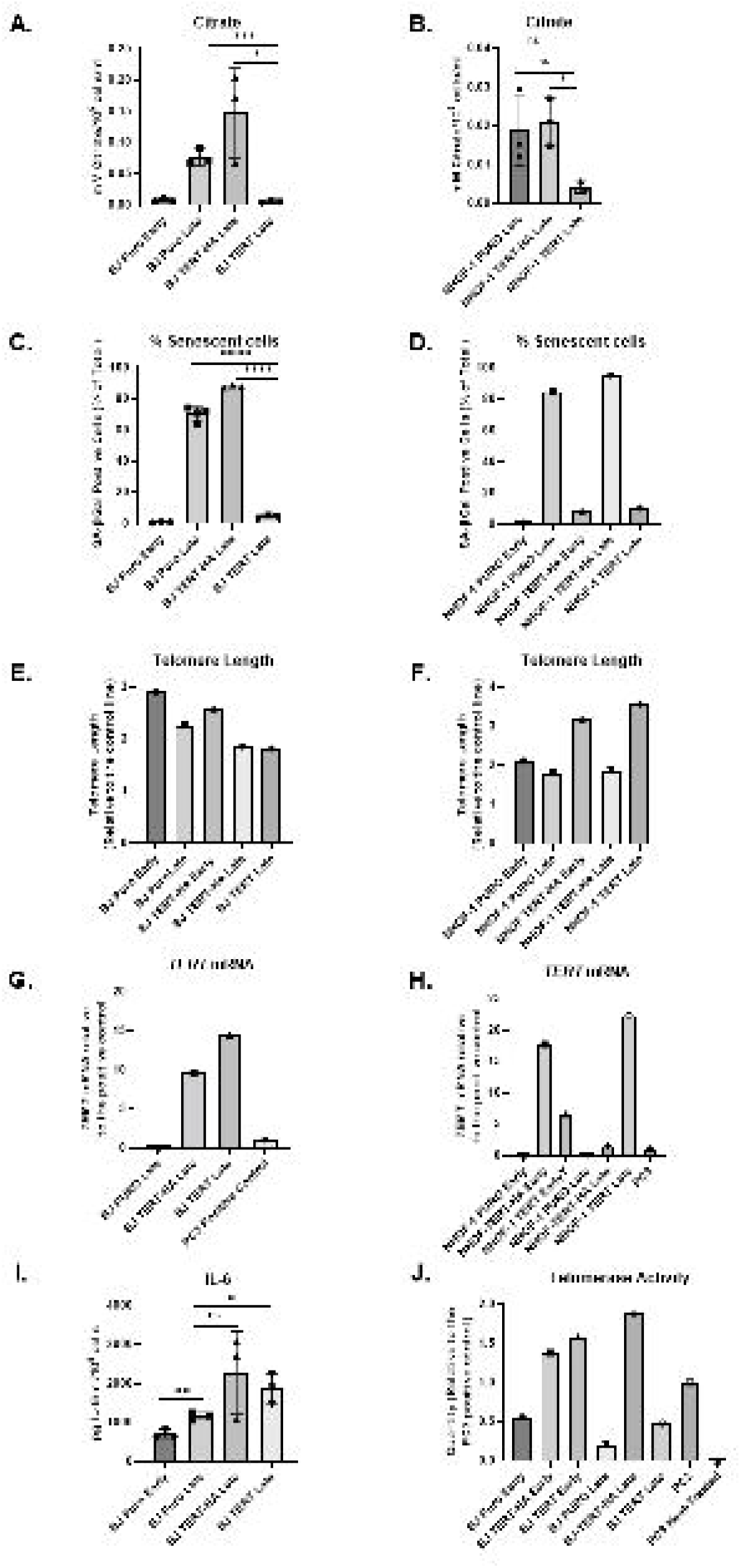
Ectopic *TERT* but not *TERT-HA* expression suppresses extracellular citrate levels in parallel with senescence bypass. **A.** and **B.** The effect ectopic *TERT* expression on citrate levels in BJ cells **(A.)** and NHOF-1 oral fibroblasts **(B.)** (n=3). Data are means +/- standard deviation. * p < 0.05; *** p < 0.001; **** p < 0.0001. **C.** and **D.** The percentage of senescent cells in **A.** and **B**. as assesed by SA-βGal **C.** n =3 and **D.** is derived from one experiment symbols in **C**. Symbols are the same as for **A.** and **B. E.** and **F.** The telomere lengths of the cells analysed in A and B. at the point of senescence showing a clear increase in telomere length in NHOF-1 cells expressing *TERT*. Data is from two replicate runs of the same samples. **G.** and **H.** *TERT* mRNA levels in the cells from **A.** and **B.** at the point of senescence. Showing high levels of *TERT* and *TERT-HA* mRNA in both BJ and NHOF-1 cells. PC3 mRNA is the postive control.**I.** IL-6 levels in the BJ cells from **A.** (n=3). Data are means +/- standard deviation. * p < 0.05; ** p < 0.01.**J.** Telomerase activity in the BJ cells from **A.** The data are derived from one experiment in **A**. PC3 is the postive control and heat-treated PC3 extract is the negative control.

### Citrate, malate and lactate are upregulated in senescent cells induced by IrrDSBs

EC is upregulated following replicative senescence of oral fibroblasts (James et al., 2015). However, IrrDSB-induced senescence (IrrDSBsen), also induces telomere dysfunction along with EC, malate and lactate and a depletion of pyruvate (all extracellular; see Supplementary Figure 1) consistent with a shift of senescent cell metabolism towards glycolysis.

### DC patients

The clinical and genetic characteristics of the DC patients are described in (Supplementary Table 1), and the age and gender of all DC and control subjects in Supplementary Table 2.

### Upregulation of plasma citrate in DC patients detected by gas chromatography/mass spectrometry (GC-MS)

First, we analysed the citrate levels DC samples (n = 28) and controls (n = 12) using the GC-MS platform. (Figure 2 top left). There was no significant difference in citrate levels when unstarved control plasma and starved serum samples were compared, suggesting that nutritional status did not affect plasma citrate levels. (Supplementary Figure 2A; P=0.60). Plasma citrate levels were resistant to haemolysis (Supplementary Figure 2B) and stable at room temperature for 24h so citrate is a highly stable plasma metabolite in healthy controls. Citrate showed a strong trend for upregulation in DC patients when compared to controls (Figure 2 top left) but the effect was of only borderline significance (P =0.06) except when patients with more severe aplastic anaemia symptoms were considered (P = 0.008). Asymptomatic DC patients in this study set were not significantly different from controls (P = 0.74).

**Figure 2.**
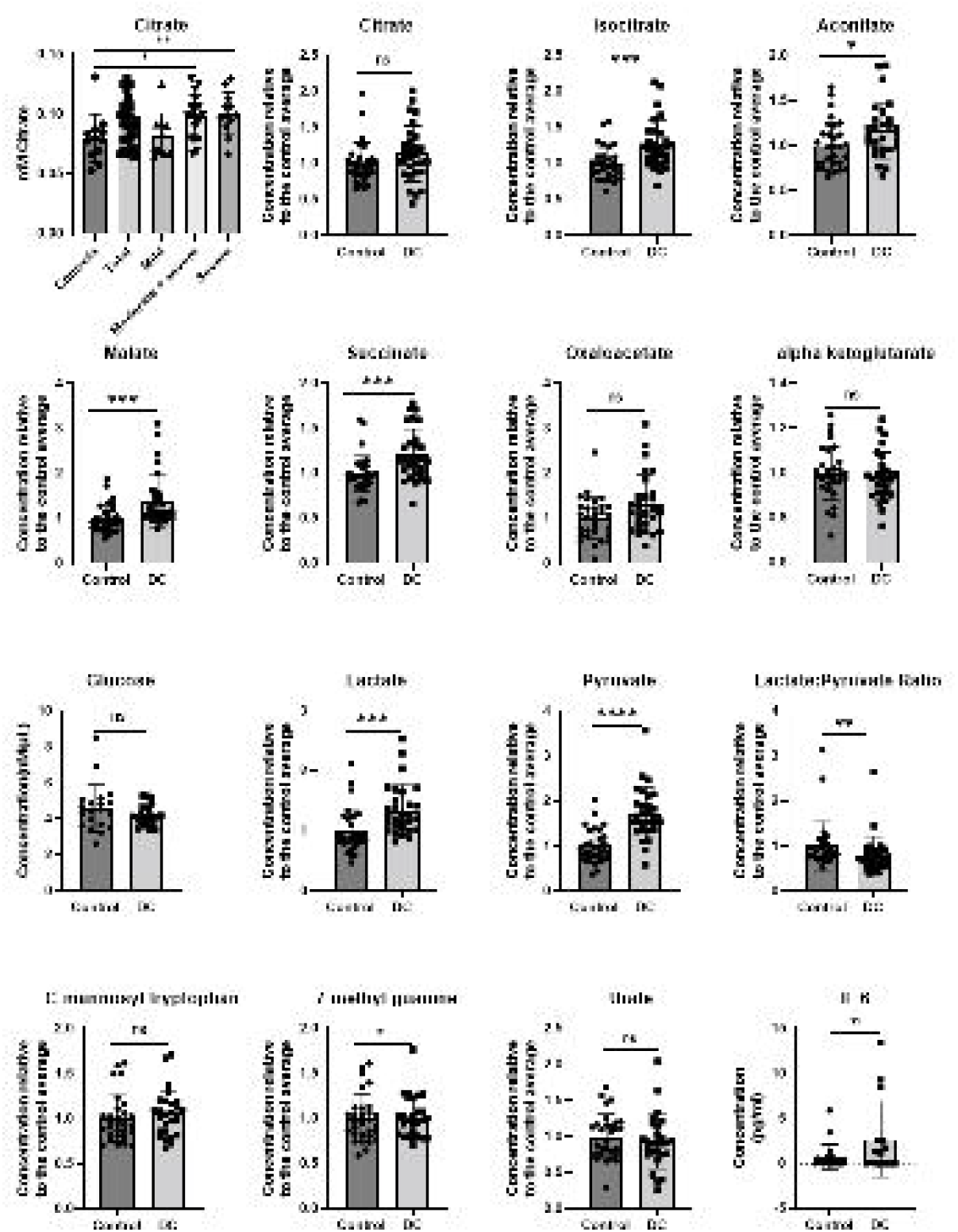
Citrate, glucose, IL-6 and normalised metabolite levels in DC patient and control subject plasma. The far left hand panel in the top row shows citrate levels in DC (n= 28) and normal control (n=12) subjects as assessed by GC-MS. DC patients with mild aplastic anaemia (0-1 abnormal clinical indicators n=10), severe (3-4 clinical indicators n=12) or a combination of severe and moderate (2-4 clinical indiators n=18) are also shown to assess the effect of aplastic anaemia on disease severity. Each point represents the average of between one and three determinations all performed at the same time. Glucose levels are in nm/µl in a subset of the DC patients (n=26) and control subjects (n=20). IL-6 levels are in pg/mL in a subset of the DC patients (n =15) and control subjects (n= 21). All remaining panels show the LC-MS metabolite levels normalised to the average of the control subject levels in DC (n=29) and control (n=30). All P values were determined by the Wilcoxon-Mann Whitney Rank Test and/or Welch’s Test and both methods gave similar P values.

### DC patients show specific changes in energy metabolism

As plasma citrate is tightly regulated *in vivo,* we considered the possibility that citrate might be further metabolised *in vivo* to other energy metabolites. We used targeted liquid chromatography/mass spectrometry (LC-MS) on an overlapping but distinct sample set to measure energy metabolites and a subset of ESM metabolites (James et al., 2015) in DC patient (n=29) and control (n =30) plasma (Figure 2; Supplementary Figure 3). The ESM metabolites were selected because they were reported to be altered in age-related diseases *in vivo* (Cheng et al., 2015; Hocher and Adamski, 2017; Huang et al., 2018; Marron et al., 2019; Menni et al., 2013; Yeri et al., 2019).

The TCA cycle metabolites isocitrate (P = 0.0007), malate (p = 0.0005), succinate (P = 0.008) and to a lesser extent oxaloacetic acid (P = 0.06) aconitate (p = 0.03) and citrate (P = 0.08) were elevated in DC samples, but other TCA cycle metabolites such as alpha ketoglutarate (AKG: P = 0.39) were not significantly altered. Lactate (P = 0.0003) and pyruvate (P = 0.0000007) levels were consistently elevated in DC patients relative to controls (Figure 2; supplementary Figure 3; supplementary Table 3A) and were significant after correction for false discovery rate. (Supplementary Figure 3B). Significantly, glucose levels were within the normal range in DC plasma and the lactate:pyruvate ratio (LPR) was lower than normal (P = 0.005) arguing against lactic acidosis and T2D.

The *DKC1* gene is X-linked and the DC patient set had a preponderance of males. However, all changes remained when only males were considered (Figure 2; Supplementary Table 3), in patient subsets with *TERC* or *TERT* mutations/variants and in the case of pyruvate, *DKC1* as well (Supplementary Table 3), arguing against a role for non-canonical functions of *TERC* and *TERT*. With the exception of succinate all changes were significant in weakly symptomatic patients (Supplementary Table 3).

### A subset of TCA metabolites, lactate and pyruvate clearly distinguish DC patients from control subjects with high specificity and selectivity

Receiver operator characteristic (ROC) curves are shown in Figure 3 and clearly show the TCA cycle metabolites isocitrate (area under the curve (AUC) = 0.76), malate (AUC = 0.77), succinate (AUC = 0.75), lactate (AUC = 0.77) gave acceptable discrimination between controls and DC; pyruvate (AUC = 0.88) gave a value that was excellent bordering on outstanding. Thus, high plasma pyruvate alone could indicate systemic telomere attrition.

**Figure 3.**
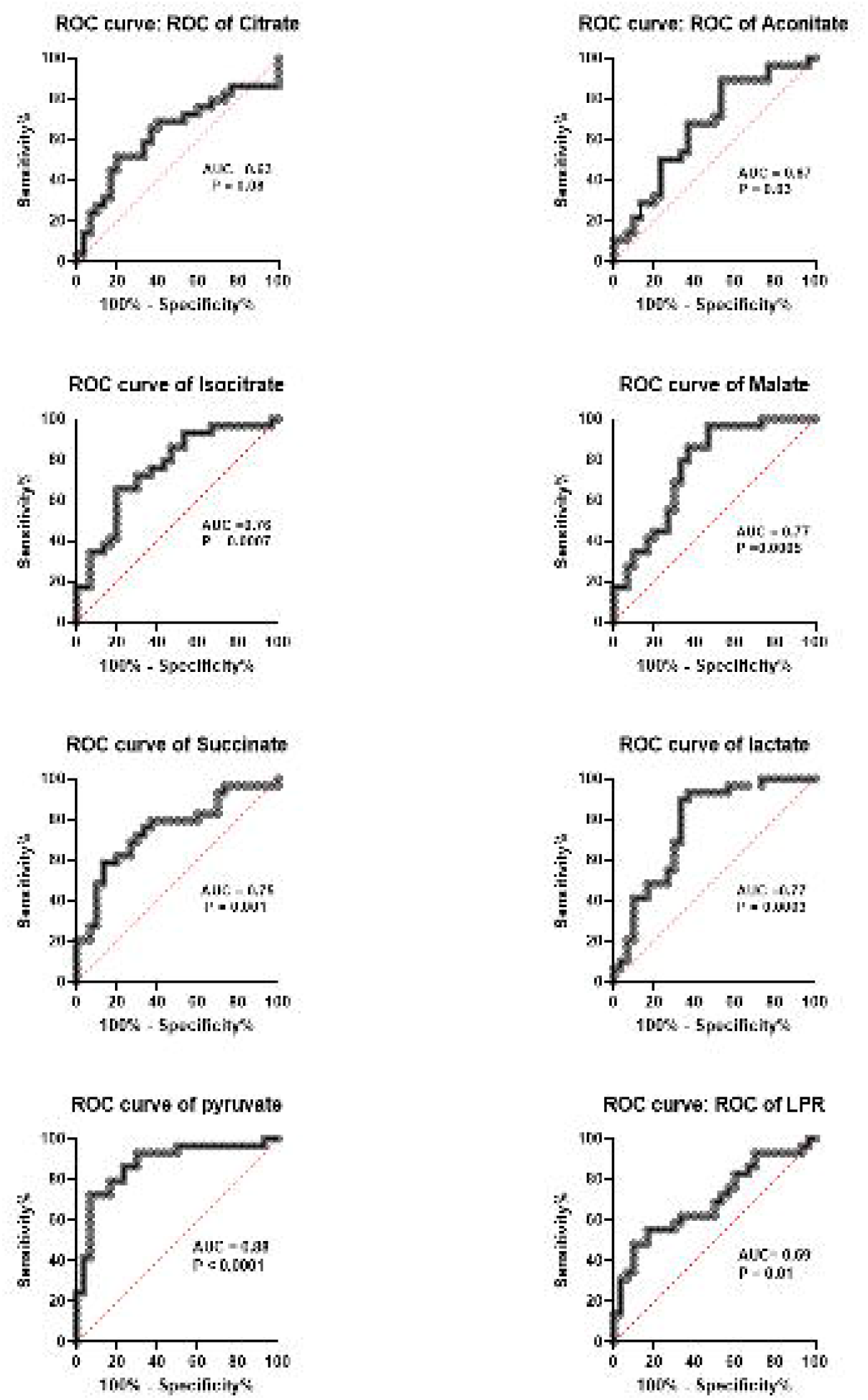
ROC curves of TCA metabolies, lactate and pyruvate in DC patient and control subject plasma. The data show ROC curves of the normalised LC/MS metabolite levels shown in Figure 2 but identical data was obtained from non-normalised values. The area under the curve (AUC) and the P values are given on each graph. An AUC of more than 0.70 is considered good and a value of more than 0.80 is considered excellent.

### Relationship of DC profile to leukocyte age-associated telomere loss (LAATL)

Linear regression analysis (Figure 4A; Supplementary Table 4) showed that only citrate (P = 0.01) and malate (0.03) levels correlated with LAATL. In the smaller GC-MS data set only citrate correlated with LAATL in the DC subgroup with *TERC* mutations by linear regression analysis (P = 0.006).

**Figure 4.**
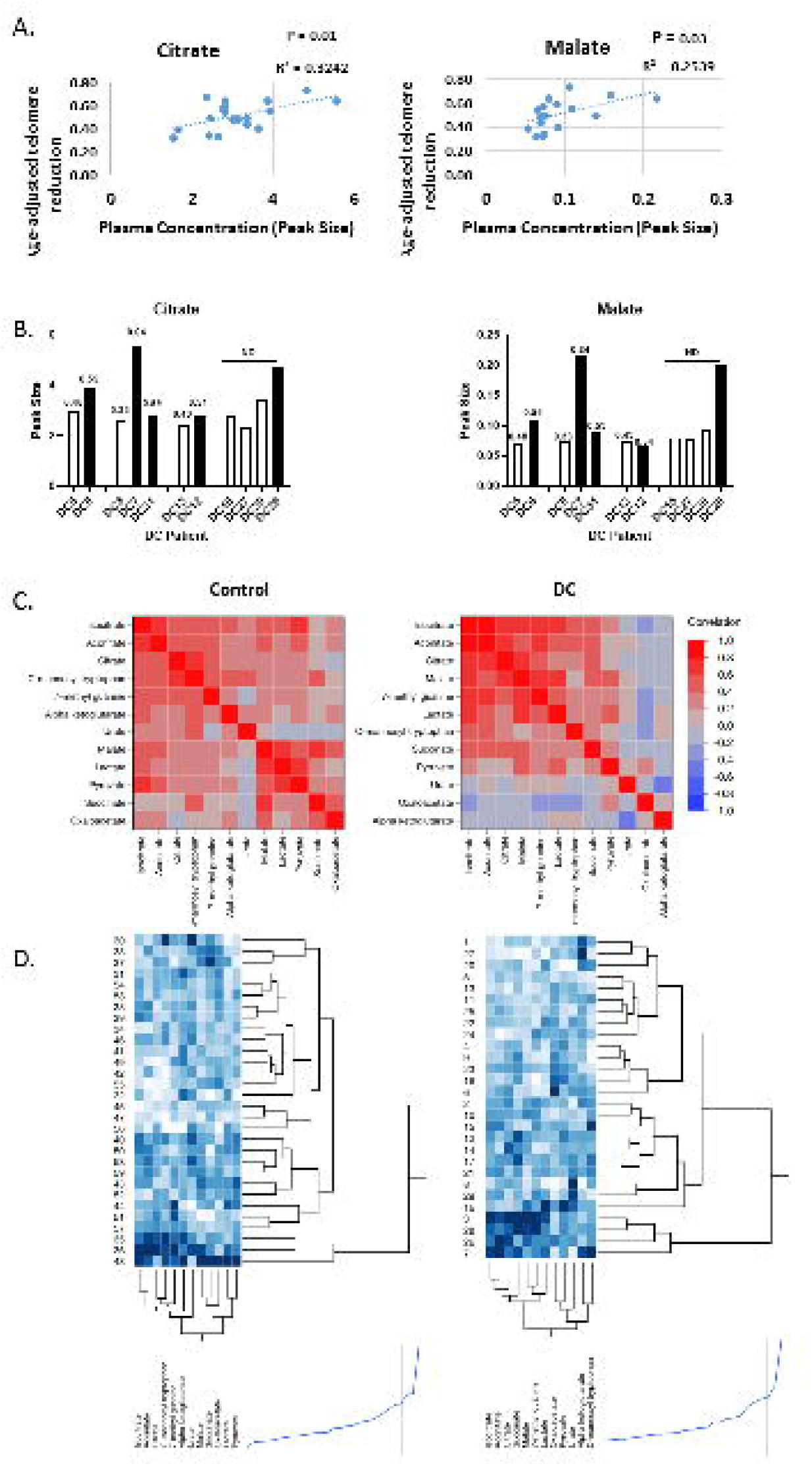
Citrate is asscoiated with DC leukocyte telomere loss and a shift in systemic energy metabolism. **A.** Linear regression analysis showing a statistically significant association between age-adjusted telomere reduction (LAATL) and plasma citrate (left panel) and malate (right panel) in DC patients (n=19). **B.** Non-normalised data from four DC familes showing that DC patients with shorter telomeres than their older relatives carrying the same mutation show a trend for increased plasma citrate (left panel) and malate (right panel). The values over the bars indicate the level of age-adjusted telomere reduction. **C.** Clustered heat map of correlations derived from normalised metabolite levels in normal subjects (left panel) and DC plasma (right panel) showing the relationship of each metabolite to citrate and showing an increased association of citrate with isocitrate, aconitate and especially malate and succinate. **D.** Cluster analysis of the data in **C.**

In addition, as with telomerase-deficient mice (Hao et al., 2005; Lee et al., 1998), LAATL increases with each generation (Vulliamy et al., 2004). We therefore examined the DC profile in individual family members from four DC families harbouring the same *TERT* or *TERC* mutation/variant. Although the numbers were small, in three families, the children had higher levels of citrate and malate than their parents/aunts did and in two families, this was associated with LAATLs and high levels of plasma IL-6 in the children. These data support the inverse relationship between these metabolites and AATL.

### DC patients show a shift in energy metabolism indicative of increased citrate catabolism

Unsupervised multivariate analyses also showed clear differences between the control and DC groups. The two groups were largely separated by principal component analysis within the first two components (Supplementary Figure 4). There was no association with patient age for the first four principal components (P > 0.05 for all). The between-metabolite correlations were also different in the two groups, as showed by both clustered heat-maps of the Pearson correlations (Figure 4C), and two-way hierarchical clustering of the two groups (Figure 4D). Unsurprisingly, the TCA metabolites citrate, isocitrate and aconitate clustered together in normal subject plasma whilst lactate, pyruvate and malate formed a separate cluster. However, in DC plasma, citrate, aconitate and isocitrate became much more strongly associated with malate and succinate indicating a shift in systemic metabolism in DC patients. In addition, isocitrate and malate became more strongly associated with lactate in DC patient plasma. Linear regression analysis (Supplementary Table 5) and scatter plot matrices (Supplementary Figures 5A and 5B) further supported these conclusions. Furthermore, supervised linear discriminant analysis (LDA) correctly classified 24 out of 30 control and 24 out of 28 DC patient samples; however as the AUC (0.90) was not significantly different from the best single metabolite AUC values (95% confidence intervals 0.87-0.99, 5000 bootstraps), we did not explore the use of LDA further.

### DC patients show elevated levels of energy metabolites in the absence of high levels of SASP and other ESM factors

We did not have access to tissue biopsies of DC patients so to estimate the amount of cellular senescence we measured several SASP and ESM factors. Interleukin 1 alpha (IL-1α) and interleukin 6 (IL-6) levels were measured in some of the samples. IL-6 levels were generally undetectable in control subjects, as expected, but were also very low in most of the DC samples and not significantly different from controls (P= 0.09). IL-1α levels were below detection limits in all but one of the DC patients (Figure 2; supplementary Figure 2; supplementary Table 3A).

In addition, we tested ESM metabolites, previously associated with ageing (Menni et al., 2013), mortality (Hocher and Adamski, 2017; Huang et al., 2018) and age-related disease (Hocher and Adamski, 2017; Huang et al., 2018). The plasma concentrations of the ESM metabolites urate, 7-methyl guanine (7-MG) and C-mannosyl tryptophan were not significantly different from controls (Figure 2; supplementary Figure 2; supplementary Table 3). The data indicate a low level of cellular senescence in DC patients.

### DC profile and SASP factors

Only citrate (P = 0.01 when analysed by GC-MS), isocitrate (P = 0.02), and 7-MG (P = 0.008) correlated with plasma IL-6 levels (Supplementary Table 6), indicating that the energy metabolite changes were independent of IL-6.

### DC profile and clinical parameters, age and gender

In both the GC-MS and LC/MS data sets there was no significant relationship between any of the above changes with any clinical indicators of aplastic anaemia, gender, donor age or different control batches and with the exception of oxaloacetic acid and urate repeat samples where available varied by less than 25% (1-22%). (Supplementary Tables 7-10). In aplastic anaemia plasma TCA metabolites are reported to be low (Zhong et al., 2015) so it is unlikely our data are the consequence of disease. Low dose Danazol treatment in a small patient subset, had no effect on the results, succinate excepted (Supplementary Table 11).

## Discussion

Telomere dysfunction is associated with senescence in human disease (Bilgili et al., 2019; Birch et al., 2016; Nault et al., 2019) but its influence on human metabolism and in particular the plasma biomarkers of age-related diseases, has not been investigated.

We report here a plasma metabolic signature of human telomere dysfunction where, despite a low predicted senescent cell burden, some TCA metabolites, lactate and pyruvate were consistently elevated in DC patient plasma but with a low LPR. The ESM metabolites citrate and malate correlated with LAATL but downstream metabolites of citrate were better at separating DC patients from controls, indicating further, or tissue specific, metabolism of citrate. As citrate is regulated by the canonical function of telomerase *in vitro* independently of the canonical SASP cytokine, IL-6, this suggests that EC accumulation is regulated independently from IL-6 and is not merely a consequence of inflammation. DC plasma metabolites were not associated with any marker of aplastic anaemia, nor any particular disease phenotype, and is unlikely to be a consequence of disease. Trivial explanations such as cell lysis and exosome release are inconsistent with the normal levels of the cytoplasmic metabolites urate and AKG in DC plasma. The low LPR and normal levels of urate distinguish the DC metabolic profile from T2D, most respiratory chain disorders and all forms of acidosis (Robinson, 2006; Shaham et al., 2010; Thompson Legault et al., 2015).

Significantly, virtually all aspects of the DC profile increase with chronological age and age-related disease (Auro et al., 2014; Cheng et al., 2015; Marron et al., 2019; Menni et al., 2013; Mota-Martorell et al., 2021; Yeri et al., 2019). High isocitrate, aconitate and malate are associated with frailty, cardiovascular disease and mortality in humans (Cheng et al., 2015; Marron et al., 2019; Yeri et al., 2019). Therefore, the plasma energy alterations are not limited to DC. AKG, which extends lifespan and protects against frailty and inflammaging in mice (Asadi Shahmirzadi et al., 2020), was not significantly altered and low plasma levels of this metabolite have not been reported in recent studies of aged or frail humans (Auro et al., 2014; Cheng et al., 2015; Marron et al., 2019; Mota-Martorell et al., 2021). However, plasma AKG increases in centenarians (Mota-Martorell et al., 2021) and although this may reflect further telomere attrition or age-related disease, it could reflect an adaptive mechanism in long-lived individuals.

Although it is difficult to speculate on mechanistic details from plasma metabolite levels the resemblance to pyruvate dehydrogenase complex (PDHC) deficiency (Thompson Legault et al., 2015) is consistent with a shift towards glycolysis, previously reported in senescent human fibroblasts *in vitro* (James et al., 2015). Notably, the high levels of isocitrate and aconitate in DC plasma are opposite to those in humans deficient in the ROS-sensitive (Lushchak et al., 2014) mitochondrial aconitase (ACO2) (Abela et al., 2017), arguing against an increase in mitochondrial ROS in DC patients and consequently, mild telomere attrition in dividing cells.

Alterations in TCA metabolites have not been reported in telomerase-deficient mice (Tomas-Loba et al., 2013) but the genetic (Hao et al., 2005) and dietary (Sun et al., 2020) differences between the mouse models and DC as well as the metabolomic platforms employed may partially explain this.

Only citrate and malate levels correlated with LAATL yet isocitrate, succinate, lactate and pyruvate were consistently elevated in DC plasma. Telomere attrition may have different direct consequences on metabolism in other cell types such as the liver.

An alternative hypothesis is that elevated plasma citrate in DC patients enters the liver and kidneys to drive the TCA cycle to cause altered metabolism. A recent study conducted in pigs offers some support for this hypothesis. Sampling of arterial and venous blood in pigs showed that citrate taken up by the kidneys, presumably via the NaDC1 and NaDC3 transporters (Mycielska et al., 2009), contributes to the TCA cycle to produce malate and succinate (Jang et al., 2019), which accumulate in kidney tissue (Jang et al., 2019). This could explain the higher plasma levels of these and other TCA metabolites in DC plasma and with chronological age.

Lactate and pyruvate levels are also associated with chronological ageing in humans (Auro et al., 2014; Menni et al., 2013; Mota-Martorell et al., 2021). Lactate and pyruvate metabolism takes place predominantly in the liver and kidneys (Jang et al., 2019) and increased plasma pyruvate was the most consistent change we observed in DC patients. Increased plasma lactate from senescent cells is oxidised by the liver and kidneys to pyruvate (Jang et al., 2019) to regulate lactate levels and could produce the low lactate:pyruvate ratio seen in the DC patient plasma.

Regardless of the underlying mechanism, the systemic accumulation of energy metabolites in DC and ageing subjects would likely accelerate ageing and cellular senescence as caloric restriction does the reverse (Fontana et al., 2018).

The most important aspect of the data described here is that a subset of plasma metabolites (isocitrate, malate, succinate, lactate and pyruvate) outperformed SASP factors in distinguishing DC patients from controls. Such metabolites may be valuable indicators of telomere dysfunction in humans and consequently therapies designed to reverse it (Bernardes de Jesus et al., 2011; Bernardes de Jesus et al., 2012; Dow and Harley, 2016).

In summary, although mechanistic details are still to be elucidated, we provide evidence that plasma metabolomics may have considerable utility in the monitoring of telomere dysfunction in human disease, in regenerative medicine and anti-ageing therapies. In addition, combining metabolomics with defined human mutations may shed light on the underlying mechanisms of ageing and contribute to the early diagnosis of age-related diseases.

## STAR METHODS

### RESOURCE AVAILABILITY

This study did not generate new unique reagents.

#### Lead contact

Further information and requests for resources and reagents should be directed to and will be fulfilled by the lead contact, Eric Kenneth Parkinson.

#### Materials availability

This study did not generate new unique reagents.

#### Data and code availability

- The Metabolon Inc. data reported in this study cannot be deposited in a public repository, because as metabolomics databases do not accept Metabolon Inc. data. All raw data files for data obtained by Metabolon Inc. will be made available on request
- Local law prohibits depositing raw [standardized datatype] datasets derived from human samples outside of the country of origin.
- All data reported in this paper will be shared by the lead contact upon request. Code
- This paper does not report original code.

### KEY RESOURCES

#### General chemicals, culture media and Reagents

The metabolite standards (Mass Spectrometry Metabolite Library of Standards, MSMLS) were purchased from IROA Technologies (Michigan, MI, U.S.A.).

Acetic acid, tributylamine, L-phenyl-^2^H_5_-alanine, 7-methylguanosine, cis-aconitic acid and α-C-mannosyltryptophan were obtained from Sigma-Aldrich (Gillingham, U.K.).

LC-MS grade water, water with 0.1% formic acid (v/v) and acetonitrile with 0.1% formic acid (v/v) were purchased from Fisher Scientific (Leicester, U.K.).

Methanol and isopropanol were obtained from Honeywell (Charlotte, NC, U.S.A.). AccQTag Ultra reagent was from Waters Corporation (Milford, MA, U.S.A.).

L-Malic acid-^13^C_4_ [CLM-8065], Succinic acid-^13^C_4_ [CLM-1571], Sodium-Lactate-^13^C_3_ [CLM-1579], Alpha-ketoglutaric acid, disodium salt (1,2,3,4-^13^C_4_) [CLM-4442] and citric acid (1,5,6-carboxyl-^13^C_3_) [CLM-9876] were obtained from CK Isotopes (Desford, U.K.).

#### Cell culture, reagents and kits

Senescence detection kit was from Biovision K-320-250 (VWR Lutterworth, Leicestershire, UK).

Dulbecco’s Modified Eagles Medium with sodium pyruvate (DMEM 4.5g/L glucose Lonza (BE12-604F/12 Slough UK and Thermo Scientific # 41966029).

HyClone Foetal Clone II fetal bovine serum (Thermo Scientific SH30066.02) L-glutamine (Life Technologies 25030-081).

Penicillin and streptomycin antibiotics (Life Technologies Cat# 15070-063).

Incubator was a humid Eppendorf Galaxy S incubator maintained at 37°C with an atmosphere of 10%CO_2_/90% air.

Quantikine® ELISA Immunoassay kits were from (R&D systems, Abingdon, Oxfordshire, UK IL-6 cat# D6050 and IL-1α cat# DLA50)

Glucose Assay Kit, Abcam, Cambridge, MA.

### EXPERIMENTAL MODEL AND SUBJECT DETAILS

#### Cells

BJ cells derived from human neonatal foreskin (Bodnar et al., 1998) at 16 mean population doublings (MPDs) were a generous gift from Professor Woodring Wright of Southwestern University, Houston, Texas, USA and the normal human oral fibroblast line NHOF-1 was derived from explant cultures of normal oral mucosa (Pitiyage et al., 2011) were used at 23 MPDs . The gender of BJ cells is male and that of NHOF-1 is not currently known.

#### Mycoplasma testing

NHOF-1 was originally tested for mycoplasma using MycoFluor mycoplasma detection kit and found to be negative. Conditioned media both BJ and NHOF-1 cell line panels were tested for mycoplasma using a Lonza MycoAlert^TM^ Mycoplasma Detection Kit and found to be negative.

#### Patients, normal subjects, plasma collection and ethics

Ethical approval for the collection of normal and DC plasma samples and the metabolomics analysis was granted by the London-City and East Research Ethics Committee (certificate number 07/Q0603/5). The details of the patients and the control subjects are given in Supplementary Tables 1 and 2. The DC patients age ranged from 9 to 70 years old (37.3 +/- 16.5 mean +/- standard deviation) and the control, subjects ranged from 23 to 92 years old (38.2+/-17.0; mean +/- standard deviation). The DC patients (n =34) were 21 males and 13 females and the control subjects (n =37) were 20 males and 14 females with 3 unknown. Not all samples were investigated by both GC-MS and LC-MS but most were. In eight DC cases two or 3 repeat samples were analyzed by LC/MS. Patient symptoms, white blood cell counts, platelet counts, hemoglobin and mean red blood cell corpuscular volume were all recorded at the time of diagnosis and in subsequent clinical assessments.

## METHOD DETAILS

### QUANTIFICATION AND STATISTICAL ANALYSIS

#### Cell culture

##### Cells

BJ cells at 16 mean population doublings were a generous gift from Professor Woodring Wright of Southwestern University, Houston, Texas, USA and the normal human oral fibroblast line NHOF-1 was derived from explant cultures of normal oral mucosa.

Cells were cultured in DMEM supplemented with penicillin and streptomycin antibiotics to a concentration of 50U/mL and L-glutamine to a concentration of 2mM, containing 10% vol/vol fetal bovine serum at 37°C in an atmosphere of 10% CO2/90% air. Flasks were kept at roughly 80% confluence, and medium was replenished every 3-4 days. Once cells became more than 80% confluent, or were needed for an experiment, they were washed once with warm (37°C) PBS containing 0.02% weight/vol EDTA before being incubated for 5 minutes with PBS containing 0.1% weight/vol trypsin and0.01% weight/vol EDTA (1mL/10cm2 dish). Following cell detachment, the trypsin was neutralized by the addition of serum containing media (3mL media for every 1mL trypsin solution), and cells were counted manually using a haemocytometer to enable calculation of the cumulative mean population doublings (cmpds). Cmpds were used throughout the study as a measure of chronological age, and were calculated using the formula: mpd = 3.32((log10cell number yield)- (log10cell number input) (Munro et al., 2001).

##### Retroviral infection of BJ and NHOF-1 cells

pBABE retroviral vectors on the puromycin-resistant backbone expressing *TERT* and *TERT-HA* (Counter et al., 1998) or the empty vector were obtained from Addgene Europe, Teddington, Middlesex, UK. Retroviral vectors were transfected into Phoenix Amphotropic 293T cells (Swift et al., 2001) to create infectious amphotropic retrovirus in 48h. The conditioned medium containing the retrovirus was filtered through a 0.45μm filter to remove Phoenix A producer cells and then incubated with sub-confluent late passage NHOF-1 (45 mean population doublings - MPDs) or BJ cells (65 MPDs) for a further 48h without polybrene as this is not recommended for the infection of normal cells. The infected and mock-infected NHOF-1 and BJ cells were then trypsinized and replated at a density of 5 x 10^5^ cells per T75 flask and cultured for a further 48-72 hours before adding 1μg/mL of puromycin as a selecting agent. The puromycin was included in the medium until all the cells on the mock-infected plates had died and removed thereafter as selecting agents can reduce the replicative lifespan of normal cells (Morgan et al., 1987). Re-introducing selection before analysis did not change the expression of the *TERT* transgenes. Cells were expanded in control medium lacking puromycin, cryopreserved and serially passaged when confluent at 1.5-3 x 103 cells per cm^2^ until the control cells senesced. Medium was changed twice weekly.

#### Induction of senescence by ionizing radiation

Fibroblasts were irradiated in suspension with γ rays from a Cs source at a dose rate of 1.4Gy/min with 20 Gy to induce irreparable DNA double strand breaks, as described (James et al., 2015), and the cells were left between 0 and 20 days in culture before analysis.

#### Characterization of senescent cells

The cellular senescence status of the cells was confirmed through senescence-associated beta galactosidase (SA-βGal) activity using a commercial kit (Biovision K-320-250) which was measured again at the time at which the experiments were conducted. Briefly, the kit was brought to room temperature for 20 minutes. Following this, the cells were washed twice with calcium and magnesium-free phosphate buffered saline (PBS) fixed for 10 minutes and washed twice more before adding the SA-βGal reagent. The plates were then sealed with parafilm to prevent dehydration, covered in foil to protect from light and incubated at 37°C in a hot room or hotbox in an atmosphere of air (no CO_2_. Late passage IMR90 cells were used as a positive control.

#### Collection of conditioned medium

Conditioned medium was collected after 24 hours from the cells, centrifuged at 800 x g for 2 minutes, the supernatant removed and centrifuged again at 13,000 rpm for 2 minutes and the final supernatant snap frozen on an ethanol-dry ice bath for 15 minutes before storage at -80°C. Unconditioned medium was also prepared identically.

#### Telomere length measurement in leukocytes

Whole blood telomere lengths were measured using the monochrome multiplex quantitative PCR method (modified from (Cawthon, 2009) on a LightCycler 480 real time thermocycler (Roche). Briefly, in each well, amplification of telomeric DNA (T) and a single copy gene (S) were quantified against standard curves obtained from dilution of a reference DNA sample. The T/S ratio, obtained in triplicate for each sample, is proportional to the telomere length. This ratio was normalized to the T/S ratio of a second reference sample that was run on every plate to give a relative T/S ratio.

#### Telomere length measurement in cells

For the absolute telomere length analysis, a qPCR method performed as described in (O’Callaghan and Fenech, 2011). Briefly, genomic DNA (gDNA) was isolated and purified from samples with a QIAamp® DNA Mini Kit (Qiagen, NL). gDNA samples were then diluted to 5ng/μL per reaction. Primer sequences were as described in (O’Callaghan and Fenech, 2011). These standards primers were HPLC purified (Sigma Aldrich, US) and resuspended to a 100μM stock in molecular grade water. Reaction mixtures for each target gene, telomere or *36B4* consisted of: 10μL *Power*Up^TM^ SYBR® Green Master mix (Applied Biosystems^TM^, US), 1μL telomere forward primer (10μM) and 1μL telomere reverse primer (10μM). In some experiments the reaction mixture consisted of, 1μL 36B4 forward primer (10μM), 1μL 36B4 reverse primer (10μM), 4μL of 5ng/μL gDNA and 4μL nuclease-free water (Ambion, Invitrogen ^TM^, US) to a total volume of 20μL. Mixtures were loaded into 96 well microplate (MicroAmp ABI, US). The samples were heated at 95°C for 10 min, followed by 40 PCR cycles of 95°C for 15 sec and 60°C for 90 sec, and was analyzed with a QuantStudio 7 Flex Real-Time PCR system (Applied Biosystems^TM^, US). Each PCR plate contained two standard curves (10^3^-10^7^ serial dilutions), telomere standard and *36B4* standard, which measures telomere sequence per sample in kb and measure diploid genome copies per samples, respectively. A plasmid gDNA (pBR322) was add to each standard with a final concentration of 20ng DNA per reaction, and all samples were analyzed in triplicated including negative template controls for each standard were included in each run. The T/S ratio, obtained in triplicate for each sample, is proportional to the telomere length. This ratio was normalized to the T/S ratio of a second reference sample that was run on every plate to give a relative T/S ratio. Data analysis was performed using Prism (Graph-pad).

#### Real time measurement of telomerase activity

Telomerase activity using qPCR was performed with a QuantStudio 7 Flex Real-Time PCR system using the TRAP protocol as described in (Wege *et al*., 2003). Briefly, pellets containing 1×10^6^ cells were washed in cold 1X PBS (Gibco^TM^, US), lysed and resuspended in 150μL of TRAPeze® 1X CHAPS Lysis Buffer (Chemicon®, Merck Millipore). Protein concentration of the samples was determined using the CB-X Protein assay (G-Biosciences) according manufacture’s guidelines. Primer sequences were as described in (Kim and Wu, 1997). For the qPCR-TRAP reaction, each sample consisted of: 12,5μL *Power* SYBR® Green Master mix (Applied Biosystems^TM^, US), TS primer (0.1μg, 100ng/μL; Invitrogen ^TM^, US), ACX primer (0.05μg, 100ng/μL; Invitrogen ^TM^, US), 250ng of sample protein extract and nuclease-free water (Ambion, Invitrogen ^TM^, US) to bring a total volume of 25μL reaction then loaded into MicroAmp Optical 96 well plate (Applied Biosystems^TM^, US). The PCR plate included a positive control (PC3-hTERT), a negative control (PC3-hTERT protein boiled for 10 minutes to deactivate the telomerase enzyme) and a single standard curve, PC-3-hTERT protein extract for serially diluted ranging from 50ng-500ng. The TRAP PCR cycle started with an initial incubation at 25°C for 20 min and amplification at 95°C for 10 min, followed by 35 PCR cycles at 95°C for 30 sec and 60°C for 90 sec. PC-3-hTERT standard curve RQ values were used to calculate the telomerase activity of the unknown samples (RQ) and then results were converted to percentage and plotted in a bar graph using Prism.

#### *hTERT* mRNA analysis

Extraction of RNA from cell lines/strains was carried out using TRI Reagent (Merck Life Sciences, Dorset UK) according to the manufacturer’s instructions. Samples were then treated with Deoxyribonuclease I (DNAse I, Invitrogen; Thermo Fisher Scientific, Inc.). To remove contaminating DNA from the RNA. 1ug of DNAse 1 treated RNA was reverse-transcribed into first-strand cDNA using the High Capacity cDNA Reverse Transcription kit (Applied Biosystems; Thermo Fisher Scientific, Inc.). The primer sequences and thermal cycling parameters used to quantify fully spliced *hTERT* (primer set E4-E5) and GAPDH (primer set E1-E3) was the same as described previously (Ducrest et al., 2001) using *Power* SYBR® Green Master mix (Applied Biosystems^TM^, US). RQ data analysis was initially performed using the SDS 2.4 software (Applied Biosystems; Thermo Fisher Scientific, Inc.) then plotted as bar charts in Prism.

#### Patients, normal subjects, plasma collection and ethics

Ethical approval for the collection of normal and DC plasma samples and the metabolomics analysis was granted by the London-City and East Research Ethics Committee (certificate number 07/Q0603/5). Blood (2-10mL) was collected by venipuncture using a 19-gauge butterfly needle into EDTA vacutainers from non-starved healthy volunteers and DC patients. Within 2 hours, blood was centrifuged at 1300g for 15 minutes at 4°C to pellet the blood cells. Following this, the plasma supernatant was transferred to a 1.5 mL Eppendorf centrifuge tube and centrifuged again at 20,600g for 2 minutes at 4°C. Plasma was then transferred to a fresh Eppendorf centrifuge tube and snap frozen on a dry ice/ethanol bath for at least 15 minutes before storage at -80°C.

#### Unbiased metabolomic screen

Metabolon Inc. carried out the unbiased metabolomics analysis. The details of the metabolomics analysis have been published previously. These include sample preparation, instrumentation, conditions for mass spectrometry (liquid chromatography/tandem mass spectrometry in positive and negative ion modes, and gas chromatography/mass spectrometry), peak data reduction, and assignment of peaks to known chemical entities by comparison to metabolite library entries of purified standards, has been previously described (James et al., 2016; James and Parkinson, 2015).

Briefly, for analysis, the median signal intensity of a given biochemical was determined across all sample groups. This median was subsequently used to scale all individual samples to a median of one for the group. A minimum value was assigned when a biochemical was not detected in an individual sample (this was rare). This data is graphically presented as scaled intensity and is thus a measure of the relative level of each metabolite. For further details, see references (James et al., 2016; James and Parkinson, 2015).

#### Targeted measurement of extracellular citrate by gas chromatography/mass spectroscopy (GC-MS)

Deuterated citrate (Citrate-d_4_) was added to each sample to a final concentration of 0.1mM as an internal standard. Metabolites were then extracted using cold methanol before being dried under vacuum desiccation. The samples were re-suspended in anhydrous pyridine containing the derivitisation agents methoxyamine hydrochloride followed by N-Methyl-N-trimethylsilyltrifluoroacetamide with 1% 2,2,2-Trifluoro-N-methyl-N-(trimethylsilyl)-acetamide and Chlorotrimethylsilane (MSTFA+1%TMCS). GCMS was performed in pulsed splitless mode on a Hewlett Packard HP6890 series GC system with Agilent 6890 series injector and a 30m x 250µm capillary column (Agilent, model number 19091s-433HP5MS) using a flow rate of 1mL/minute, and a Hewlett Packard 5973 mass selective detector. The acquisition was conducted in selective ion monitoring mode, with the m/z values 273, 347, 375 and 465 for citrate, and 276, 350, 378 and 469 for citrate-d_4_. The dwell time for all of these ions was 50ms.

#### Metabolite extraction and targeted analysis by liquid chromatography/mass spectroscopy

For the analysis of organic acids from plasma, plasma samples stored at -80C were thawed at room temperature and treated with methanol to deproteinize the samples in a volumetric ratio of 4:1. The sample order was randomized before extraction. The samples were treated in polypropylene microcentrifuge tubes. The amount of plasma was 100 µl, or, for samples where this amount was not available, less, with the amount of methanol used adjusted proportionally. The samples were vortexed briefly and then kept on ice for at least 60 min. They were then centrifuged at 7378 g for 10 min at 4 °C. Multiple aliquots of the supernatant were collected and transferred to new microcentrifuge tubes. For pooled QCs sample 75 µl of each sample supernatant was mixed in a new tube. Supernatants were dried in a centrifugal vacuum concentrator (Savant Speedvac, Thermo). Dried samples were kept at -80 °C until analyses.

Samples were reconstituted for analysis in 75 µl of a solution containing stable isotope labelled (SIL) internal standards, made up in LC-MS grade water. The solution contained L-malic acid-^13^C_4_, succinic acid-^13^C_4_, sodium lactate-^13^C_3_, citric acid (1,5,6-carboxyl-^13^C_3_) and L-phenyl-d_5_-alanine (this last used to monitor the consistency of injection volumes, but not otherwise used to normalize the data). Stock solutions of each SIL organic acid were prepared at 1 mg/mL in water or methanol and stored at -20 °C until required. These were then diluted 1:1000 to produce a final working solution with concentrations of each SIL standard at 1 µg/mL. The reconstituted samples were then centrifuged and transferred to LC-MS vials.

Analysis was performed with a Waters XEVO TQ-S tandem mass spectrometer coupled with a Waters Acquity UPLC binary solvent manager equipped with a CTC autosampler. Data were acquired with electrospray ionization in negative mode, with only [M-H]^-^ ions considered. Liquid chromatography was performed using a Waters HSS T3 column (1.8 µm, 2.1 x 100 mm) with a binary solvent system of 10mM of tributylamine and 15mM of acetic acid in water (solvent A), and 80% methanol and 20% isopropanol (solvent B) with a constant flow rate of 0.4 mL/min. Column temperature was 45C. The gradient elution profile was: 0-0.5 min at 0% B; 0.5-6 min linear ramp from 0% B to 5%B; 4-6 min at 5%B; 6-6.5 min linear ramp to 20% B; 6.5-8.5 min at 20% B; 8.5-14 min linear ramp to 55%B; 14-15 min linear ramp to 100% B; 15-17min at 100% B; 17-18 min a linear ramp to 0% B; and 3 min at initial conditions before next injection cycle starts. Samples in the autosampler were held at 4 °C, and 10 µl injected to a 5µl sample loop. Sample run order was randomized to minimize batch effects. Before analysis, injections of double blanks (water) and single blanks (SIL standard mix) were performed to ensure system stability, and to determine carryover and contaminants originating from the system and from the sample preparation procedure, vials, solvents, pipettes etc. A pooled QC sample was injected at the beginning of the run and then once every 10-sample injections throughout the run, to assess instrument stability throughout the entire batch.

#### Compound specific mass spectrometry parameters

Compound specific mass spectrometry parameters were determined individually and described in (Perin et al., 2021); two transitions were monitored for each analyte to confirm its identity where possible.

The MRM conditions (parent ion → fragment ion(s); collision energy(s)) of metabolites were as follows. Pyruvate (m/z 87 → m/z 43; 8); Lactate (m/z 89 → m/z 43; 11); Lactate-^13^C_3_ (m/z 92 → m/z 45; 11); Succinate (m/z 117 → m/z 73,99; 10,9); Succinate^13^C_4_ (m/z 121 → m/z 76; 10); Oxaloacetate (m/z 131→ m/z 87; 13); Malate (m/z 133→ m/z 115, 71; 9,13); Malate-^13^C_4_ (m/z 137→ m/z 119; 9); Oxoglutarate (m/z 145→ m/z 101; 11); Oxoglutarate (1,2,3,4-^13^C_4_) (m/z 149→ m/z 105; 11); 7-Methyl Guanosine (m/z 164→ m/z 106, 121); 20,12); Urate (m/z 167→ m/z 124, 69; 14, 16); L-phenyl-^2^H_5_-alanine (m/z 169→ m/z 72, 108, 152; 18, 16, 12); Aconitate (m/z 173→ m/z 129, 111, 85; 13,22, 19); Isocitrate (m/z 191→ m/z 73, 111; 22, 12); Citrate (m/z 191→ m/z 43, 87; 19, 18); Citrate (1,5,6-carboxyl-^13^C_3_) (m/z 194.02→m/z 88, 113; 18, 12); α-C-Mannosyltryptophan (m/z 365→ m/z 245, 217; 16, 22).^a^Sodium-Lactate-^13^C_3_; ^b^Succinic acid-^13^C_4_; ^c^L-Malic acid-^13^C_4_; ^d^Alpha-ketoglutaric acid, disodium salt (1,2,3,4-^13^C_4_); ^e^L-phenyl-^2^H_5_-alanine; ^f^citric acid (1,5,6-carboxyl-^13^C_3_).

The raw LC-MS data were analyzed using Skyline (MacCoss Lab, (Pino et al., 2020). Blanks and QC samples were used to exclude contamination and low quality samples, with acceptance criteria of <0.3 relative standard deviation (RSD) for QC samples, and <1% peak area (of QC) in blanks. Metabolite concentrations were expressed as the ratio relative to the values for the SIL internal standards for each metabolite, except for those metabolites without a SIL standard available (pyruvate, isocitrate, α-C-mannosyltryptophan, urate, 7-methylguanosine, oxaloacetate, aconitate), for which relative count data were used.

#### Glucose measurement in plasma samples

Glucose concentrations in plasma were measured with an enzymatic kit specific for glucose (Glucose Assay Kit, Abcam, Cambridge, MA) following manufacturers protocol. Absorbance was measured on a SPECTROstar Nanospectrometer (BMG Labtech).

#### Interleukin 1α and interleukin 6 measurement in culture media and plasma samples

Quantikine® ELISA Immunoassay kits for human IL-1α and IL-6 (R&D Systems, Abingdon, UK) kits utilizing the sandwich ELISA method were performed to quantitate the protein concentrations closely following the manufacturer’s protocol. The detection limit for IL-1 α was 3.9 – 250 pg/mL and for IL-6 3.13-300pg/mL.

All reagents were brought to room temperature before use. All standards, samples and controls were diluted if required in the appropriate calibration buffer (plasma or culture media) and assayed in duplicate. The required microplate strips were fixed to the plate frame. All ELISAs were repeated three times from three different independent experiments.

Briefly, standards, samples and controls were added into each well pre-coated with capture antibody. With the IL-1α assay, diluent was added to each well prior to adding the samples. The plate was covered with the adhesive strip provided and incubated for 2 hours at room temperature. Any unbound antigen-antibody in each well was aspirated and washed by filling each well with 350µl wash buffer using a squirt bottle. The process was repeated twice for three washes. After the last wash, any remaining wash buffer was removed by aspirating. The plate was inverted and blotted against clean paper towels. Next, the protein conjugate was added to each well. The plate was covered with a new adhesive strip and incubated at room temperature for 1 to 2 hours. The washing steps were repeated as above to remove any non-binding antigen-antibody. Following this, the substrate solution was added into each well and incubated at room temperature for 20 minutes to produce the chromogenic reaction. This reaction is light sensitive therefore the plate was protected from light. Finally, 50µl of stop solution was added to each well to stop the reaction.

The optical density (OD) of each well was determined within 30 minutes using a Fluostar-OPTIMA microplate reader (BMG-Labtech, Aylesbury, UK) at 450nm and correction wavelength at 570nm. The 570nm correction wavelength was to correct for optical imperfections in the plate. The duplicate readings were averaged for each standard, sample and control and the averaged zero standard OD was subtracted from these readings. A log/log standard curve (the best-fit curve) was created using Excel software with the log mean absorbance for each standard on the y-axis plotted against the log concentration on the x-axis. Wherever the samples had been diluted, the concentration read from the standard curve was multiplied by the dilution factor.

#### Principal component analysis

This was carried out using autoscaled and mean-centred data. The first four PCs were inspected for associations with disease status.

#### Clustered heatmaps of correlations

Correlation matrices (calculated for the control and DC groups separately) are shown as heat maps, with correlation intensity given by the color scale.

#### Cluster analysis

Two-way hierarchical clustering was performed using standardized data and Ward’s method of linkages for the control and DC groups separately. The heat map represents the compound concentrations.

#### Supervised linear discriminant analysis (LDA)

Unsupervised and supervised multivariate analyses were carried out using principal components analysis (using autoscaled data), and Fisher’s linear discriminant analysis, respectively”. If you think it needs mentioning, the software used was JMP Pro 15.

#### False discovery rates (FDRs)

FDRs were determined using Graphpad prism and adaptive linear step-up procedures that control the FDR.

#### Other statistical methods

Cell culture and unbiased metabolomics screen data and was analyzed by the Student’s unpaired T test. Raw and normalized plasma metabolomics data was analyzed by the Wilcoxon-Mann-Whitney and Mann-Whitney U tests and corrected in the latter case for false discovery rate. Linear regression analysis was conducted by using the Excel data analysis package and the graphs prepared in Excel.

## Supplemental items

**Supplementary Figure 1.** Changes in extracellular metabolites following IrrDSB-induced senescence.

**Supplementary Figure 2.** Citrate Levels are simialr in starved and unstarved subjects and resistant to haemolysis.

**Supplementary Figure 3.** Non-normalized plasma metabolite levels in control subjects and DC patients as assessed by LC-MS

**Supplementary Figure 4.** Principal Component Analysis (PCA) of cluster metabolomics.

**Supplementary Figure 5.** Scatter plot matrices.

**Table 1.** Clinical and Genetic Characteristics of DC Patients.

**Table 2.** Age and Gender of control subjects.

**Table 3A.** Rank Test P values of DC Patients (n=29) versus normal donors (n =30)

**Table 3B.** Rank Test P values of DC Patients (n=29) versus normal donors (n =30) normalized to control average and corrected for false discovery rate.

**Table 4.** Linear regression P values of plasma levels of metabolites versus LAATL in DC patients.

**Table 5.** Relationship of Citrate to other metabolites by linear regression analysis.

**Table 6.** Linear regression P values of plasma levels of metabolites versus plasma. Levels of IL-6 in *DC* patients (n = 15 by GC-MS and LC-MS).

**Table 7.** Linear regression P values of plasma levels of metabolites versus DC clinical indicators of aplastic anemia in DC patients.

**Table 8.** The effect of age and gender on LC-MS metabolites.

**Table 9.** Effect of sample time and batch on metabolites in normal subjects

**Table 10.** Average variation between repeat samples of eight DC patients.

**Table 11.** The effect of low dose Danazol treatment in DC patients (n =6) v no treatment (n =21).

## Acknowledgments

The authors would like to thank the Dunhill Medical Trust (grant number R452/1115) for the financial support of Emma James. Karen-Ng Lee Peng received a Ph.D. scholarship (Hadiah Latihan Persekutuan) from the Malaysian Ministry of Education. Professor Tom Vulliamy for performing the telomere length analysis of the human leukocytes, Dr. Harriet Allan and Professor Tim Warner for assistance with blood samples from normal volunteers and Dr. Amy Lewis for technical advice. We would also like to thank Dr. Fabian Flores-Borja for valuable discussions regarding the metabolism of citrate and Professors Andrew Silver and Julie Adam (University of Oxford) for critical reading of the manuscript.

## Author contributions

ENJ and MB performed the Gas Chromatography/Mass Spectroscopy analysis of the conditioned medium and plasma samples; V. S-K performed the Liquid Chromatography/Mass Spectroscopy analysis of plasma samples; M.M., E.K.P and J.B. performed the statistical analysis of all the data; E.K.P and J.B. prepared the figures; K-N. LP. performed the ELISA assays; T.R. performed the qPCR analysis of *TERT* and telomerase on the fibroblast lines; S.M. performed the telomere length analysis. I.D. provided the patient samples, the telomere length data and all the clinical data in addition to obtaining ethical approval for the study; E.K.P conceived the study, obtained funding, performed most of the *in vitro* experiments and wrote the initial draft of the manuscript; E.K.P. M.M. I.D and J.B analysed the data and finalised the writing of the manuscript.

---

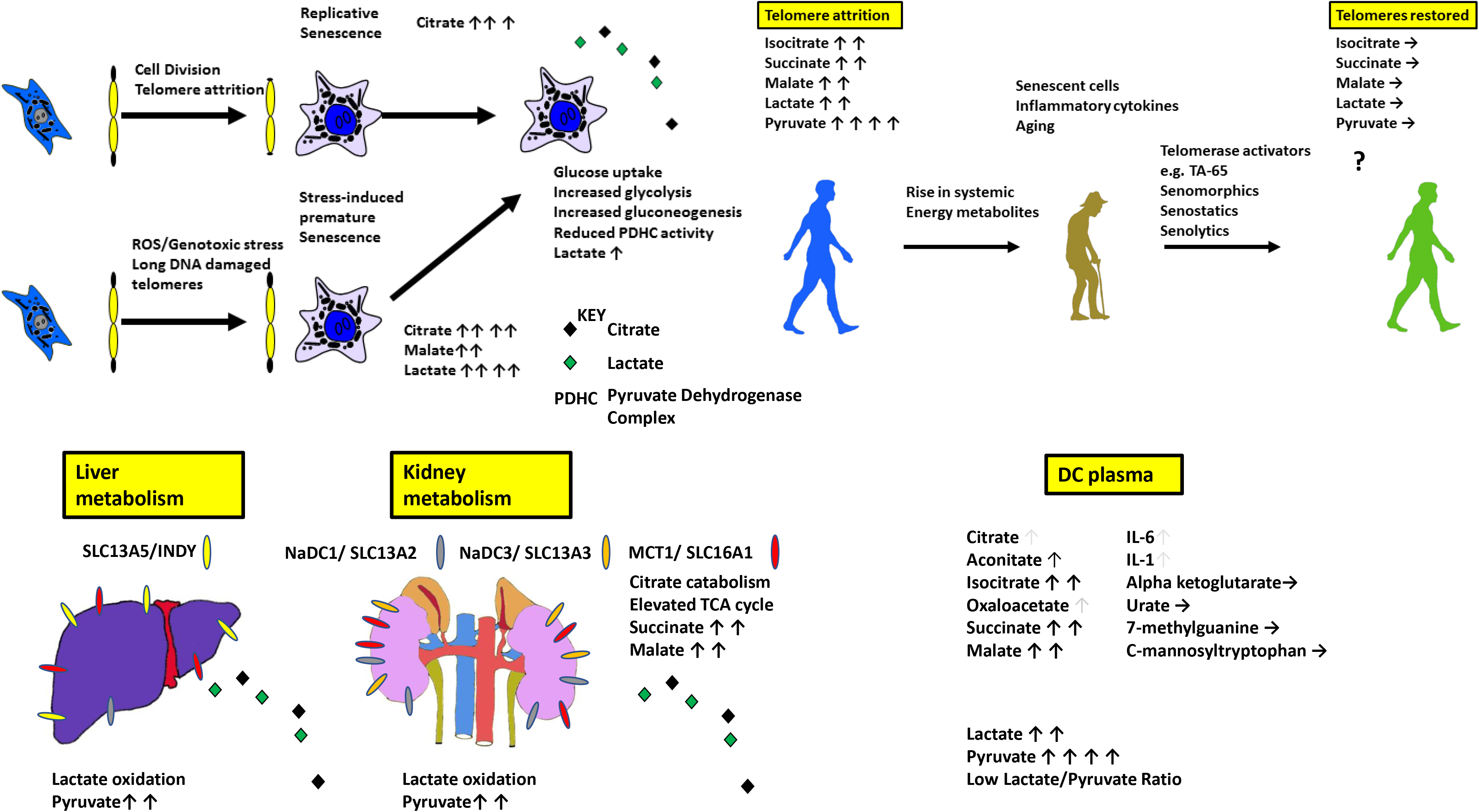

## References

1. Abela, L., Spiegel, R., Crowther, L.M., Klein, A., Steindl, K., Papuc, S.M., Joset, P., Zehavi, Y., Rauch, A., Plecko, B., et al. (2017). Plasma metabolomics reveals a diagnostic metabolic fingerprint for mitochondrial aconitase (ACO2) deficiency. PLoS One 12, e0176363.

2. Asadi Shahmirzadi, A., Edgar, D., Liao, C.Y., Hsu, Y.M., Lucanic, M., Asadi Shahmirzadi, A., Wiley, C.D., Gan, G., Kim, D.E., Kasler, H.G., et al. (2020). Alpha-Ketoglutarate, an Endogenous Metabolite, Extends Lifespan and Compresses Morbidity in Aging Mice. Cell Metab 32, 447–456 e446.

3. Auro, K., Joensuu, A., Fischer, K., Kettunen, J., Salo, P., Mattsson, H., Niironen, M., Kaprio, J., Eriksson, J.G., Lehtimaki, T., et al. (2014). A metabolic view on menopause and ageing. Nat Commun 5, 4708.

4. Bernardes de Jesus, B., Schneeberger, K., Vera, E., Tejera, A., Harley, C.B., and Blasco, M.A. (2011). The telomerase activator TA-65 elongates short telomeres and increases health span of adult/old mice without increasing cancer incidence. Aging Cell 10, 604–621.

5. Bernardes de Jesus, B., Vera, E., Schneeberger, K., Tejera, A.M., Ayuso, E., Bosch, F., and Blasco, M.A. (2012). Telomerase gene therapy in adult and old mice delays aging and increases longevity without increasing cancer. EMBO Mol Med 4, 691–704.

6. Bilgili, H., Bialas, A.J., Gorski, P., and Piotrowski, W.J. (2019). Telomere Abnormalities in the Pathobiology of Idiopathic Pulmonary Fibrosis. J Clin Med 8.

7. Birch, J., Victorelli, S., Rahmatika, D., Anderson, R.K., Jiwa, K., Moisey, E., Ward, C., Fisher, A.J., De Soyza, A., and Passos, J.F. (2016). Telomere Dysfunction and Senescence-associated Pathways in Bronchiectasis. Am J Respir Crit Care Med 193, 929–932.

8. Birkenfeld, A.L., Lee, H.Y., Guebre-Egziabher, F., Alves, T.C., Jurczak, M.J., Jornayvaz, F.R., Zhang, D., Hsiao, J.J., Martin-Montalvo, A., Fischer-Rosinsky, A., et al. (2011). Deletion of the mammalian INDY homolog mimics aspects of dietary restriction and protects against adiposity and insulin resistance in mice. Cell Metab 14, 184–195.

9. Bodnar, A.G., Ouellette, M., Frolkis, M., Holt, S.E., Chiu, C.P., Morin, G.B., Harley, C.B., Shay, J.W., Lichtsteiner, S., and Wright, W.E. (1998). Extension of life-span by introduction of telomerase into normal human cells. Science 279, 349–352.

10. Brachs, S., Winkel, A.F., Tang, H., Birkenfeld, A.L., Brunner, B., Jahn-Hofmann, K., Margerie, D., Ruetten, H., Schmoll, D., and Spranger, J. (2016). Inhibition of citrate cotransporter Slc13a5/mINDY by RNAi improves hepatic insulin sensitivity and prevents diet-induced non-alcoholic fatty liver disease in mice. Mol Metab 5, 1072–1082.

11. Campisi, J., Kapahi, P., Lithgow, G.J., Melov, S., Newman, J.C., and Verdin, E. (2019). From discoveries in ageing research to therapeutics for healthy ageing. Nature 571, 183–192.

12. Cawthon, R.M. (2009). Telomere length measurement by a novel monochrome multiplex quantitative PCR method. Nucleic Acids Res 37, e21.

13. Cheng, S., Larson, M.G., McCabe, E.L., Murabito, J.M., Rhee, E.P., Ho, J.E., Jacques, P.F., Ghorbani, A., Magnusson, M., Souza, A.L., et al. (2015). Distinct metabolomic signatures are associated with longevity in humans. Nat Commun 6, 6791.

14. Coppe, J.P., Patil, C.K., Rodier, F., Sun, Y., Munoz, D.P., Goldstein, J., Nelson, P.S., Desprez, P.Y., and Campisi, J. (2008). Senescence-associated secretory phenotypes reveal cell-nonautonomous functions of oncogenic RAS and the p53 tumor suppressor. PLoS Biol 6, 2853–2868.

15. Counter, C.M., Hahn, W.C., Wei, W., Caddle, S.D., Beijersbergen, R.L., Lansdorp, P.M., Sedivy, J.M., and Weinberg, R.A. (1998). Dissociation among in vitro telomerase activity, telomere maintenance, and cellular immortalization. Proc Natl Acad Sci U S A 95, 14723–14728.

16. d’Adda di Fagagna, F., Reaper, P.M., Clay-Farrace, L., Fiegler, H., Carr, P., Von Zglinicki, T., Saretzki, G., Carter, N.P., and Jackson, S.P. (2003). A DNA damage checkpoint response in telomere-initiated senescence. Nature 426, 194–198.

17. Dokal, I. (2011). Dyskeratosis congenita. Hematology Am Soc Hematol Educ Program 2011, 480–486.

18. Dow, C.T., and Harley, C.B. (2016). Evaluation of an oral telomerase activator for early age-related macular degeneration - a pilot study. Clin Ophthalmol 10, 243–249.

19. Ducrest, A.L., Amacker, M., Mathieu, Y.D., Cuthbert, A.P., Trott, D.A., Newbold, R.F., Nabholz, M., and Lingner, J. (2001). Regulation of human telomerase activity: repression by normal chromosome 3 abolishes nuclear telomerase reverse transcriptase transcripts but does not affect c-Myc activity. Cancer Res 61, 7594–7602.

20. Epel, E.S., Blackburn, E.H., Lin, J., Dhabhar, F.S., Adler, N.E., Morrow, J.D., and Cawthon, R.M. (2004). Accelerated telomere shortening in response to life stress. Proc Natl Acad Sci U S A 101, 17312–17315.

21. Fan, S.Z., Sung, C.W., Tsai, Y.H., Yeh, S.R., Lin, W.S., and Wang, P.Y. (2021). Nervous System Deletion of Mammalian INDY in Mice Mimics Dietary Restriction-Induced Memory Enhancement. J Gerontol A Biol Sci Med Sci 76, 50–56.

22. Fontana, L., Nehme, J., and Demaria, M. (2018). Caloric restriction and cellular senescence. Mech Ageing Dev 176, 19–23.

23. Fumagalli, M., Rossiello, F., Clerici, M., Barozzi, S., Cittaro, D., Kaplunov, J.M., Bucci, G., Dobreva, M., Matti, V., Beausejour, C.M., et al. (2012). Telomeric DNA damage is irreparable and causes persistent DNA-damage-response activation. Nat Cell Biol 14, 355–365.

24. Gadalla, S.M., Cawthon, R., Giri, N., Alter, B.P., and Savage, S.A. (2010). Telomere length in blood, buccal cells, and fibroblasts from patients with inherited bone marrow failure syndromes. Aging (Albany NY) 2, 867–874.

25. Guo, N., Parry, E.M., Li, L.S., Kembou, F., Lauder, N., Hussain, M.A., Berggren, P.O., and Armanios, M. (2011). Short telomeres compromise beta-cell signaling and survival. PLoS One 6, e17858.

26. Hao, L.Y., Armanios, M., Strong, M.A., Karim, B., Feldser, D.M., Huso, D., and Greider, C.W. (2005). Short telomeres, even in the presence of telomerase, limit tissue renewal capacity. Cell 123, 1121–1131.

27. Hastie, N.D., Dempster, M., Dunlop, M.G., Thompson, A.M., Green, D.K., and Allshire, R.C. (1990). Telomere reduction in human colorectal carcinoma and with ageing. Nature 346, 866–868.

28. Hewitt, G., Jurk, D., Marques, F.D., Correia-Melo, C., Hardy, T., Gackowska, A., Anderson, R., Taschuk, M., Mann, J., and Passos, J.F. (2012). Telomeres are favoured targets of a persistent DNA damage response in ageing and stress-induced senescence. Nat Commun 3, 708.

29. Hickson, L.J., Langhi Prata, L.G.P., Bobart, S.A., Evans, T.K., Giorgadze, N., Hashmi, S.K., Herrmann, S.M., Jensen, M.D., Jia, Q., Jordan, K.L., et al. (2019). Senolytics decrease senescent cells in humans: Preliminary report from a clinical trial of Dasatinib plus Quercetin in individuals with diabetic kidney disease. EBioMedicine 47, 446–456.

30. Hocher, B., and Adamski, J. (2017). Metabolomics for clinical use and research in chronic kidney disease. Nat Rev Nephrol 13, 269–284.

31. Huang, J., Weinstein, S.J., Moore, S.C., Derkach, A., Hua, X., Liao, L.M., Gu, F., Mondul, A.M., Sampson, J.N., and Albanes, D. (2018). Serum Metabolomic Profiling of All-Cause Mortality: A Prospective Analysis in the Alpha-Tocopherol, Beta-Carotene Cancer Prevention (ATBC) Study Cohort. Am J Epidemiol 187, 1721–1732.

32. James, E.L., Lane, J.A., Michalek, R.D., Karoly, E.D., and Parkinson, E.K. (2016). Replicatively senescent human fibroblasts reveal a distinct intracellular metabolic profile with alterations in NAD+ and nicotinamide metabolism. Sci Rep 6, 38489.

33. James, E.L., Michalek, R.D., Pitiyage, G.N., de Castro, A.M., Vignola, K.S., Jones, J., Mohney, R.P., Karoly, E.D., Prime, S.S., and Parkinson, E.K. (2015). Senescent human fibroblasts show increased glycolysis and redox homeostasis with extracellular metabolomes that overlap with those of irreparable DNA damage, aging, and disease. J Proteome Res 14, 1854–1871.

34. James, E.L., and Parkinson, E.K. (2015). Serum metabolomics in animal models and human disease. Curr Opin Clin Nutr Metab Care 18, 478–483.

35. Jang, C., Hui, S., Zeng, X., Cowan, A.J., Wang, L., Chen, L., Morscher, R.J., Reyes, J., Frezza, C., Hwang, H.Y., et al. (2019). Metabolite Exchange between Mammalian Organs Quantified in Pigs. Cell Metab 30, 594–606 e593.

36. Jaskelioff, M., Muller, F.L., Paik, J.H., Thomas, E., Jiang, S., Adams, A.C., Sahin, E., Kost-Alimova, M., Protopopov, A., Cadinanos, J., et al. (2011). Telomerase reactivation reverses tissue degeneration in aged telomerase-deficient mice. Nature 469, 102–106.

37. Justice, J.N., Nambiar, A.M., Tchkonia, T., LeBrasseur, N.K., Pascual, R., Hashmi, S.K., Prata, L., Masternak, M.M., Kritchevsky, S.B., Musi, N., et al. (2019). Senolytics in idiopathic pulmonary fibrosis: Results from a first-in-human, open-label, pilot study. EBioMedicine 40, 554–563.

38. Leandro, J.G., Espindola-Netto, J.M., Vianna, M.C., Gomez, L.S., DeMaria, T.M., Marinho-Carvalho, M.M., Zancan, P., Paula Neto, H.A., and Sola-Penna, M. (2016). Exogenous citrate impairs glucose tolerance and promotes visceral adipose tissue inflammation in mice. Br J Nutr 115, 967–973.

39. Lee, H.W., Blasco, M.A., Gottlieb, G.J., Horner, J.W., 2nd, Greider, C.W., and DePinho, R.A. (1998). Essential role of mouse telomerase in highly proliferative organs. Nature 392, 569–574.

40. Lushchak, O.V., Piroddi, M., Galli, F., and Lushchak, V.I. (2014). Aconitase post-translational modification as a key in linkage between Krebs cycle, iron homeostasis, redox signaling, and metabolism of reactive oxygen species. Redox Rep 19, 8–15.

41. Marron, M.M., Harris, T.B., Boudreau, R.M., Clish, C.B., Moore, S.C., Murphy, R.A., Murthy, V.L., Sanders, J.L., Shah, R.V., Tseng, G.C., et al. (2019). Metabolites Associated with Vigor to Frailty Among Community-Dwelling Older Black Men. Metabolites 9.

42. Menni, C., Kastenmuller, G., Petersen, A.K., Bell, J.T., Psatha, M., Tsai, P.C., Gieger, C., Schulz, H., Erte, I., John, S., et al. (2013). Metabolomic markers reveal novel pathways of ageing and early development in human populations. Int J Epidemiol 42, 1111–1119.

43. Mitchell, J.R., Wood, E., and Collins, K. (1999). A telomerase component is defective in the human disease dyskeratosis congenita. Nature 402, 551–555.

44. Morgan, J.R., Barrandon, Y., Green, H., and Mulligan, R.C. (1987). Expression of an exogenous growth hormone gene by transplantable human epidermal cells. Science 237, 1476–1479.

45. Mota-Martorell, N., Jove, M., Borras, C., Berdun, R., Obis, E., Sol, J., Cabre, R., Pradas, I., Galo-Licona, J.D., Puig, J., et al. (2021). Methionine transsulfuration pathway is upregulated in long-lived humans. Free Radic Biol Med 162, 38–52.

46. Munoz-Lorente, M.A., Cano-Martin, A.C., and Blasco, M.A. (2019). Mice with hyper-long telomeres show less metabolic aging and longer lifespans. Nat Commun 10, 4723.

47. Munro, J., Steeghs, K., Morrison, V., Ireland, H., and Parkinson, E.K. (2001). Human fibroblast replicative senescence can occur in the absence of extensive cell division and short telomeres. Oncogene 20, 3541–3552.

48. Mycielska, M.E., Dettmer, K., Rummele, P., Schmidt, K., Prehn, C., Milenkovic, V.M., Jagla, W., Madej, G.M., Lantow, M., Schladt, M., et al. (2018). Extracellular Citrate Affects Critical Elements of Cancer Cell Metabolism and Supports Cancer Development In Vivo. Cancer Res 78, 2513–2523.

49. Mycielska, M.E., Milenkovic, V.M., Wetzel, C.H., Rummele, P., and Geissler, E.K. (2015). Extracellular Citrate in Health and Disease. Curr Mol Med 15, 884–891.

50. Mycielska, M.E., Patel, A., Rizaner, N., Mazurek, M.P., Keun, H., Patel, A., Ganapathy, V., and Djamgoz, M.B. (2009). Citrate transport and metabolism in mammalian cells: prostate epithelial cells and prostate cancer. Bioessays 31, 10–20.

51. Nakamura, K., Izumiyama-Shimomura, N., Sawabe, M., Arai, T., Aoyagi, Y., Fujiwara, M., Tsuchiya, E., Kobayashi, Y., Kato, M., Oshimura, M., et al. (2002). Comparative analysis of telomere lengths and erosion with age in human epidermis and lingual epithelium. J Invest Dermatol 119, 1014–1019.

52. Nault, J.C., Ningarhari, M., Rebouissou, S., and Zucman-Rossi, J. (2019). The role of telomeres and telomerase in cirrhosis and liver cancer. Nat Rev Gastroenterol Hepatol 16, 544–558.

53. Patel, P.L., Suram, A., Mirani, N., Bischof, O., and Herbig, U. (2016). Derepression of hTERT gene expression promotes escape from oncogene-induced cellular senescence. Proc Natl Acad Sci U S A 113, E5024–5033.

54. Perin, G., Fletcher, T., Sagi-Kiss, V., Gaboriau, D.C.A., Carey, M.R., Bundy, J.G., and Jones, P.R. (2021). Calm on the surface, dynamic on the inside. Molecular homeostasis of Anabaena sp. PCC 7120 nitrogen metabolism. Plant Cell Environ 44, 1885–1907.

55. Pino, L.K., Searle, B.C., Bollinger, J.G., Nunn, B., MacLean, B., and MacCoss, M.J. (2020). The Skyline ecosystem: Informatics for quantitative mass spectrometry proteomics. Mass Spectrom Rev 39, 229–244.

56. Pitiyage, G.N., Slijepcevic, P., Gabrani, A., Chianea, Y.G., Lim, K.P., Prime, S.S., Tilakaratne, W.M., Fortune, F., and Parkinson, E.K. (2011). Senescent mesenchymal cells accumulate in human fibrosis by a telomere-independent mechanism and ameliorate fibrosis through matrix metalloproteinases. J Pathol 223, 604–617.

57. Robinson, B.H. (2006). Lactic acidemia and mitochondrial disease. Mol Genet Metab 89, 3–13.

58. Rogina, B. (2017). INDY-A New Link to Metabolic Regulation in Animals and Humans. Front Genet 8, 66.

59. Sahin, E., Colla, S., Liesa, M., Moslehi, J., Muller, F.L., Guo, M., Cooper, M., Kotton, D., Fabian, A.J., Walkey, C., et al. (2011). Telomere dysfunction induces metabolic and mitochondrial compromise. Nature 470, 359–365.

60. Sellami, M., Bragazzi, N., Prince, M.S., Denham, J., and Elrayess, M. (2021). Regular, Intense Exercise Training as a Healthy Aging Lifestyle Strategy: Preventing DNA Damage, Telomere Shortening and Adverse DNA Methylation Changes Over a Lifetime. Front Genet 12, 652497.

61. Shaham, O., Slate, N.G., Goldberger, O., Xu, Q., Ramanathan, A., Souza, A.L., Clish, C.B., Sims, K.B., and Mootha, V.K. (2010). A plasma signature of human mitochondrial disease revealed through metabolic profiling of spent media from cultured muscle cells. Proc Natl Acad Sci U S A 107, 1571–1575.

62. Sun, C., Wang, K., Stock, A.J., Gong, Y., Demarest, T.G., Yang, B., Giri, N., Harrington, L., Alter, B.P., Savage, S.A., et al. (2020). Re-equilibration of imbalanced NAD metabolism ameliorates the impact of telomere dysfunction. EMBO J 39, e103420.

63. Suram, A., Kaplunov, J., Patel, P.L., Ruan, H., Cerutti, A., Boccardi, V., Fumagalli, M., Di Micco, R., Mirani, N., Gurung, R.L., et al. (2012). Oncogene-induced telomere dysfunction enforces cellular senescence in human cancer precursor lesions. EMBO J 31, 2839–2851.

64. Suvakov, S., Ghamrawi, R., Cubro, H., Tu, H., White, W.M., Tobah, Y.S.B., Milic, N.M., Grande, J.P., Cunningham, J.M., Chebib, F.T., et al. (2021). Epigenetic and senescence markers indicate an accelerated ageing-like state in women with preeclamptic pregnancies. EBioMedicine 70, 103536.

65. Swift, S., Lorens, J., Achacoso, P., and Nolan, G.P. (2001). Rapid production of retroviruses for efficient gene delivery to mammalian cells using 293T cell-based systems. Curr Protoc Immunol Chapter 10, Unit 10 17C.

66. Thompson Legault, J., Strittmatter, L., Tardif, J., Sharma, R., Tremblay-Vaillancourt, V., Aubut, C., Boucher, G., Clish, C.B., Cyr, D., Daneault, C., et al. (2015). A Metabolic Signature of Mitochondrial Dysfunction Revealed through a Monogenic Form of Leigh Syndrome. Cell Rep 13, 981–989.

67. Tomas-Loba, A., Bernardes de Jesus, B., Mato, J.M., and Blasco, M.A. (2013). A metabolic signature predicts biological age in mice. Aging Cell 12, 93–101.

68. Tomas-Loba, A., Flores, I., Fernandez-Marcos, P.J., Cayuela, M.L., Maraver, A., Tejera, A., Borras, C., Matheu, A., Klatt, P., Flores, J.M., et al. (2008). Telomerase reverse transcriptase delays aging in cancer-resistant mice. Cell 135, 609–622.

69. Vulliamy, T., Marrone, A., Szydlo, R., Walne, A., Mason, P.J., and Dokal, I. (2004). Disease anticipation is associated with progressive telomere shortening in families with dyskeratosis congenita due to mutations in TERC. Nat Genet 36, 447–449.

70. Vulliamy, T.J., Knight, S.W., Mason, P.J., and Dokal, I. (2001). Very short telomeres in the peripheral blood of patients with X-linked and autosomal dyskeratosis congenita. Blood Cells Mol Dis 27, 353–357.

71. Wang, Q., Zhan, Y., Pedersen, N.L., Fang, F., and Hagg, S. (2018). Telomere Length and All-Cause Mortality: A Meta-analysis. Ageing Res Rev 48, 11–20.

72. Wellen, K.E., Hatzivassiliou, G., Sachdeva, U.M., Bui, T.V., Cross, J.R., and Thompson, C.B. (2009). ATP-citrate lyase links cellular metabolism to histone acetylation. Science 324, 1076–1080.

73. Willmes, D.M., Daniels, M., Kurzbach, A., Lieske, S., Bechmann, N., Schumann, T., Henke, C., El-Agroudy, N.N., Da Costa Goncalves, A.C., Peitzsch, M., et al. (2021). The longevity gene mIndy (I’m Not Dead, Yet) affects blood pressure through sympathoadrenal mechanisms. JCI Insight 6.

74. Yeri, A., Murphy, R.A., Marron, M.M., Clish, C., Harris, T.B., Lewis, G.D., Newman, A.B., Murthy, V.L., and Shah, R.V. (2019). Metabolite Profiles of Healthy Aging Index Are Associated With Cardiovascular Disease in African Americans: The Health, Aging, and Body Composition Study. J Gerontol A Biol Sci Med Sci 74, 68–72.

75. Yousefzadeh, M.J., Schafer, M.J., Noren Hooten, N., Atkinson, E.J., Evans, M.K., Baker, D.J., Quarles, E.K., Robbins, P.D., Ladiges, W.C., LeBrasseur, N.K., et al. (2018). Circulating levels of monocyte chemoattractant protein-1 as a potential measure of biological age in mice and frailty in humans. Aging Cell 17.

76. Zhong, P., Zhang, J., and Cui, X. (2015). Abnormal metabolites related to bone marrow failure in aplastic anemia patients. Genet Mol Res 14, 13709–13718.

