## Supplementary material for "Telomerase deficiency in humans is associated with systemic age-related changes in energy metabolism": Supplementary Information..docx

**Supplementary Figures**
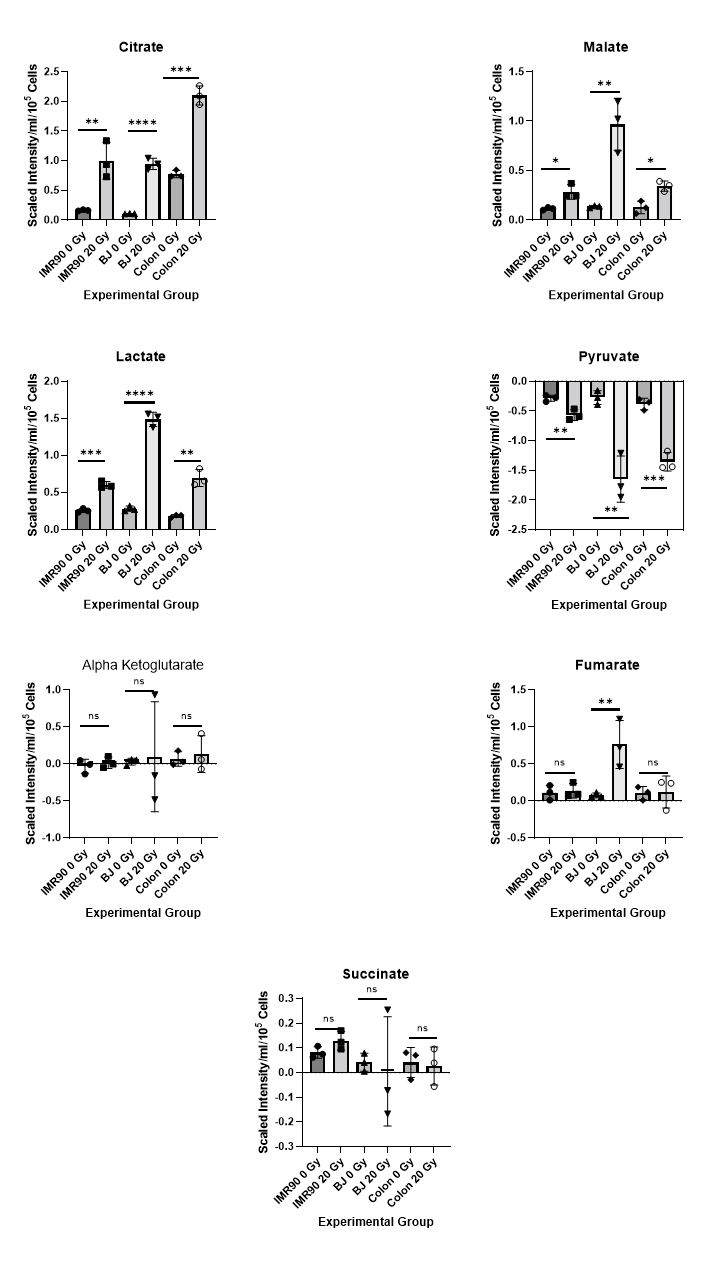

**Supplementary Figure 1 Changes in extracellular metabolites following IrrDSB-induced senescence.**

Data mined from (James et al., 2015) shows that citrate, malate and lactate are statistically elevated in 3 different IrrDSB-induced senescent fibroblast cell lines (20 Gy) relative to the non-irradiated control (0 Gy. Senescence induced by 20Gy of ᵞ irradiation followed by 20 days of incubation to yield >90% senescent cells compared to <10% in the controls. Data are means +/- standard deviation. * p < 0.05;** p < 0.01; *** p < 0.001; **** p < 0.0001 as assessed by Student’s unpaired T test n=3 for each line.

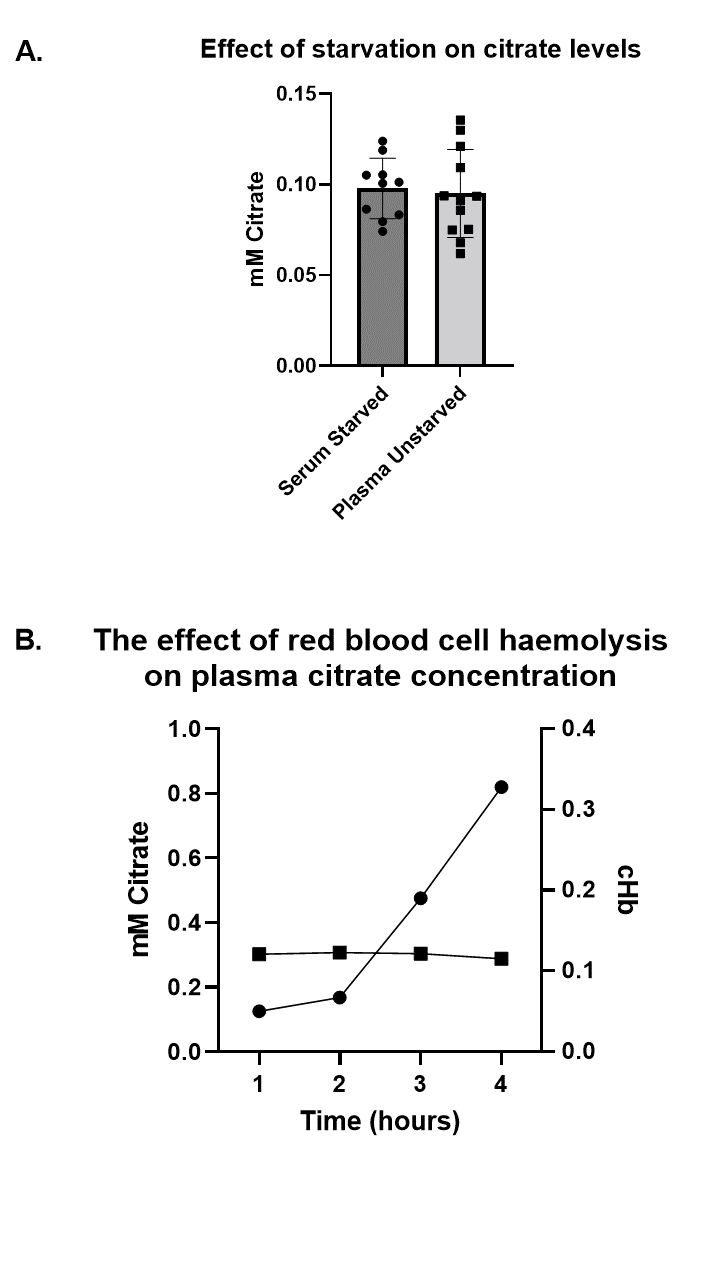

**Supplementary Figure 2**. **Citrate Levels are simialr in starved and unstarved subjects and resistant to haemolysis.**

A. Shows the comparison between citrate levels in serum from starved (n =10) and plasma from unstarved (N =13) normal subjects as measured by GC/MS. The results were not significant as assessed by the Wilcoxon-Mann-Whitney rank test and the Welch’s T test.
B. Shows the red blood cell lysis of a control blood sample followed by the measurement of haemaglobin (cHb) and the measurement of citrate by GC/MS. The experiment shows that as cHb (circles) rises over time the citrate levels (squares) do not change. The experiment shows that citrate levels are resistent to haemolysis.

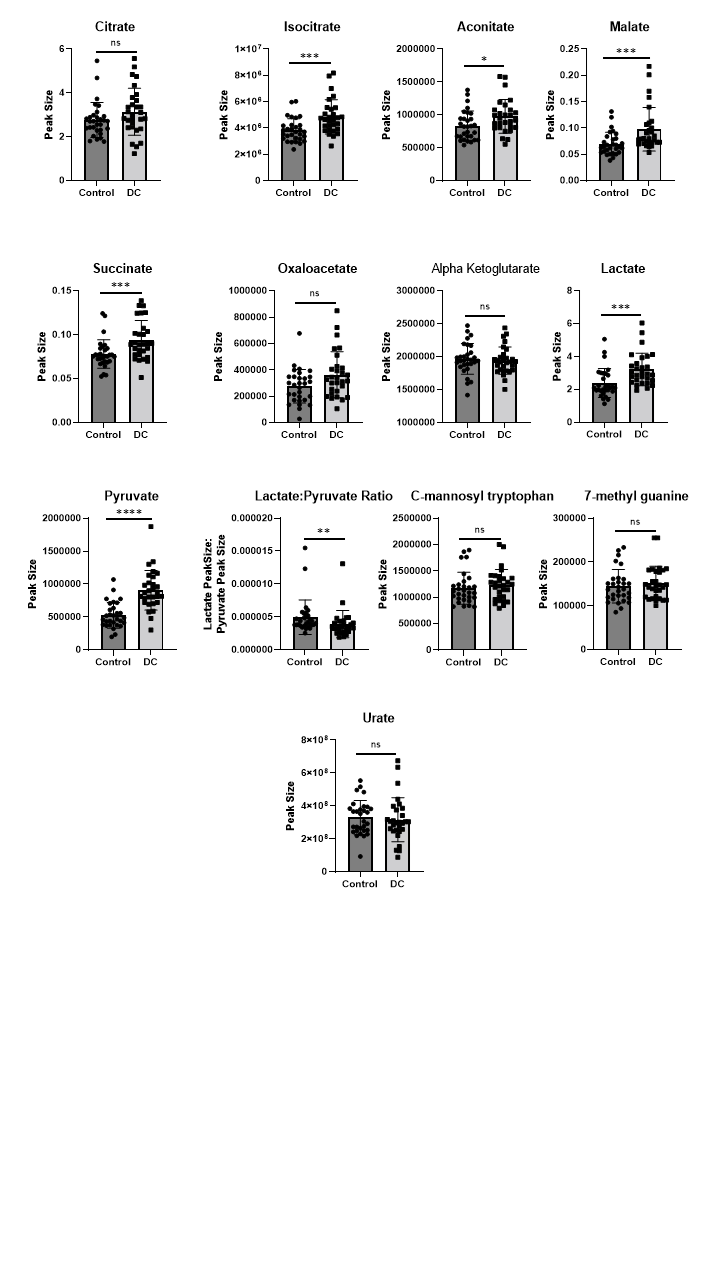

**Supplementary Figure 3. Non-normalised plasma metabolite levels in control subjects and DC patients as assessed by LC/MS**

**T**he figure shows the non-normalised data from which the data in Figures 2, 3, 4C, 4D and Supplementary Figures 4 and 5, were derived. The data shows LC/MS data for normal subjects (n = 30) and DC patients (n =29) * p < 0.05;** p < 0.01; *** p < 0.001; **** p < 0.0001 as assessed by the Wilcoxon-Mann-Whitney test.

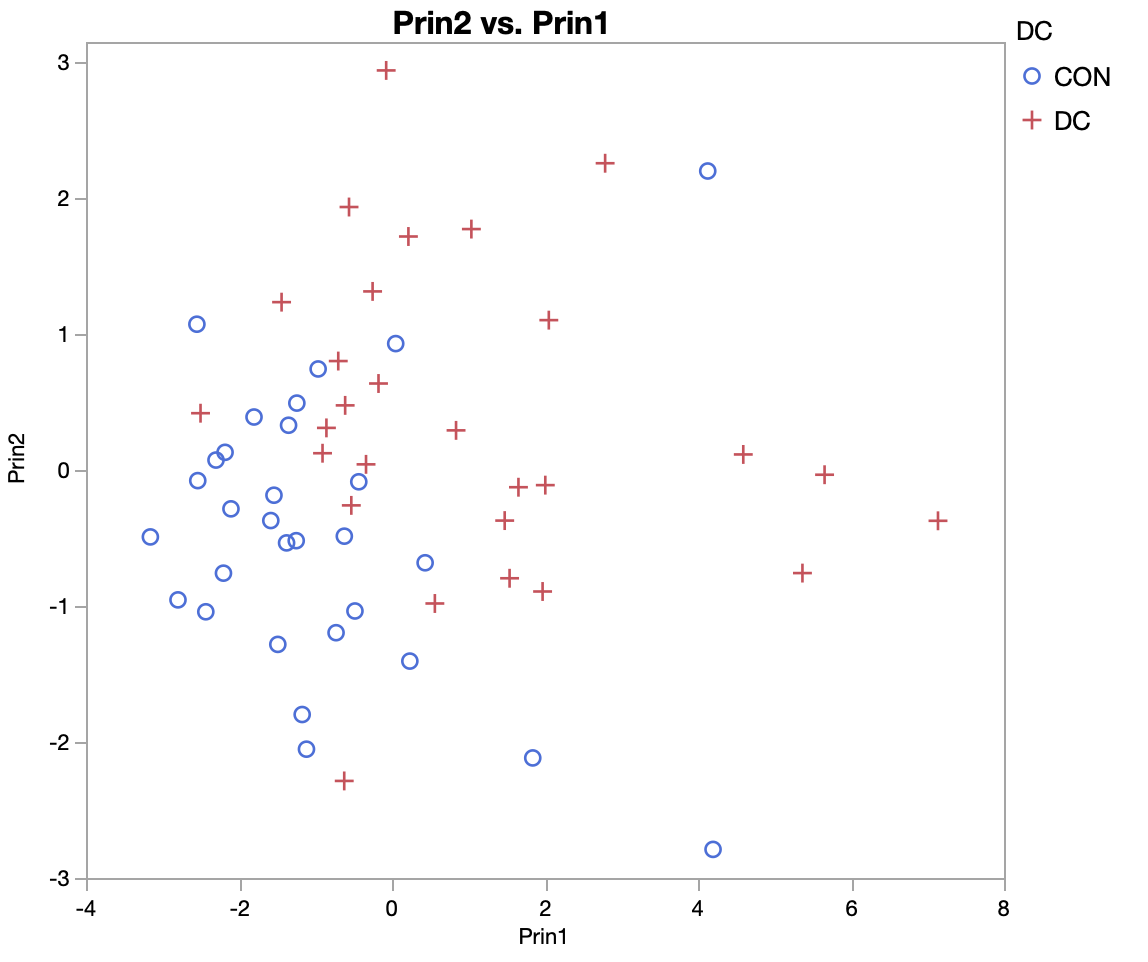

**Supplementary Figure 4. Principal Component Analysis (PCA) of metabolite concentrations demonstrates that there is a clear unsupervised difference between the groups.**

Control group: blue circles. DC group: red crosses. PCA based on autoscaled data. The group separation falls across the first two principal components; the separation between the sum values (PC 1 score + PC 2 score) is highly significant (P = 0.000006, t test; AUROC = 0.86).

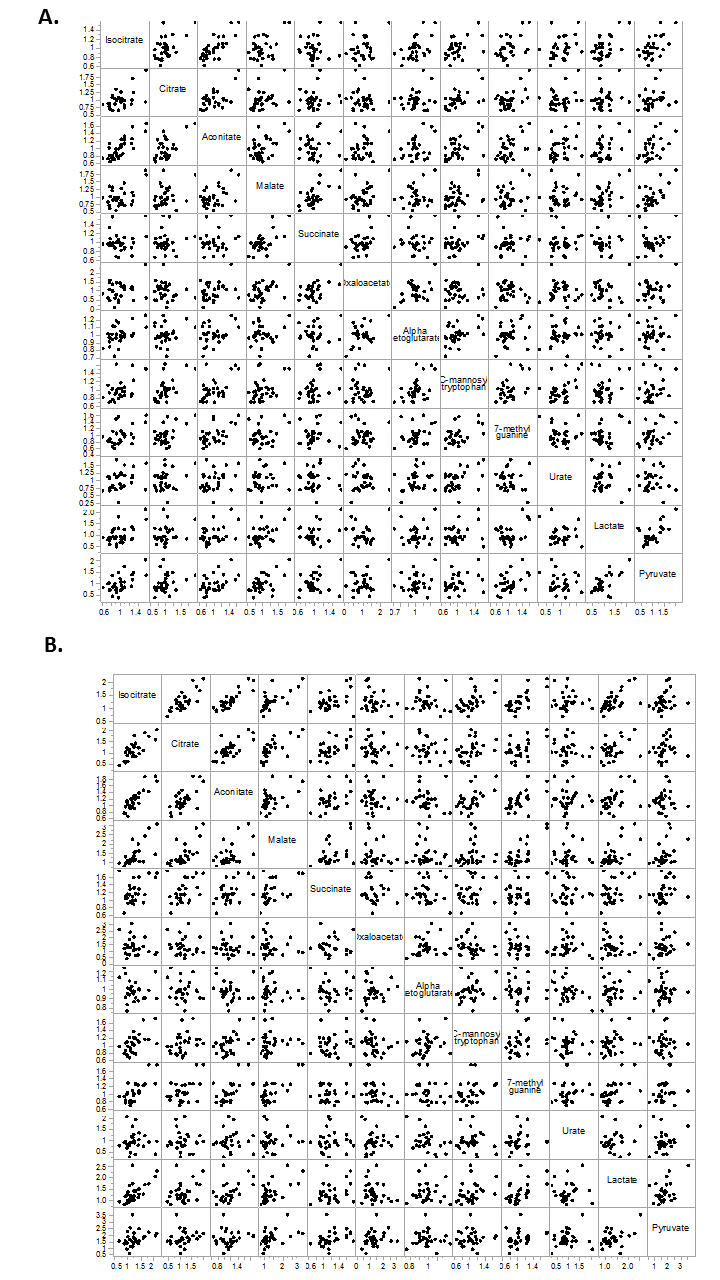

**Supplementary Figure 5 Scatter plot matrices.**

The matrices show multiple linear regression analyses summarising the relationships between the different metabolites in **A.** normal subject and **B.** DC patient plasma.

**Table 1. Clinical and Genetic Characteristics of DC Patients.**

| Patient Number | Gender | Age | Mutation/Variant | Treatment |
| --- | --- | --- | --- | --- |
| DC 1 | F | 32 | *TERT* mutation. Bone marrow failure. | None |
| DC 2 | M | 27 | *DKC1* mutation. Muco-cutaneous features and bone marrow failure. | None |
| DC 3 | M | 61 | *TERC* mutation c287C-->G. Interstitial lung disease | None |
| DC 5* | M | 67 | *TERC* mutation c180C-->T. Mild anaemia. | None |
| DC 6* | M | 39 | *TERC* mutation c180C-->T. Bone marrow failure. | None |
| DC 7+ | M | 37 | *TERC* mutation c377 A-->G. Bone marrow failure and liver disease. | Danazol 200mg twice a day |
| DC 8+ | M | 70 | *TERC* mutation c377 A-->G. Interstitial lung disease. | Danazol 100mg twice a day |
| DC 9 | M | 27 | *TERC* variant. Bone marrow failure. | None |
| DC 10 | M | 23 | *DKC1* mutation 402Gly-->Glu. Muco-cutaneous features. | None |
| DC 11 | M | 26 | *DKC1* mutation. Muco-cutaneous features. | None |
| DC 12** | M | 31 | *TERC* mutation 287C-->G. Asymptomatic. | None |
| DC 13** | F | 59 | *TERC* mutation 287C-->G . Interstitial lung disease. | Prednisolone 7.5mg/day |
| DC 14 | M | 29 | *TERT* mutation pAla716Thr. Bone marrow failure and nail dystrophy | None |
| DC 15 + | F | 45 | *TERT* variant, Ile1130Val. Bone marrow failure and liver function abnormalities. | None |
| DC 16 | F | 12 | NOP10 homozygous mutation. Muco-cutaneous features and cataracts. | None |
| DC 17 | M | 24 | *TERC* mutation. Bone marrow failure and muco-cutaneous features. | None |
| DC 18 | F | 51 | *TERT* mutation. Bone marrow failure and liver disease | Danazol 100mg twice a day |
| DC 19 | F | 29 | *TERC* mutation. Bone marrow failure and muco-cutaneous features. | Danazol 100mg twice a day |
| DC 20 | F | 38 | *TERC* mutation c180C-->T. Bone marrow failure and skin abnormalities | None |
| DC 21 | M | 52 | Heterozygous RTEL1 mutation. Liver disease and myelodysplasia. | None |
| DC 22 | M | 34 | *DKC1* mutation. Muco-cutaneous features. | None |
| DC 23 | M | 22 | *DKC1* mutation. Muco-cutaneous features. | Danazol 100mg alternate days |
| DC 24 | M | 52 | *DKC1* mutation Gly402 Glu. Muco-cutaneous features | Danazol  100mg every 3 days |
| DC 25 | F | 38 | *TERT* Variant. Bone marrow failure. | None |
| DC26 +/++ | M | 9 | *TERT* variant Ile 1130 Val. Dry skin and abnormal teeth. Son of *DC*15. | None |
| DC27++ | F | 46 | *TERT* variant Ile 1130 Val. Sibling of *DC*15. Dry skin | None |
| DC 28 | M | 17 | Uncharacterised DC. Bone marrow failure and nail dystrophy | None |
| DC 29 | F | 13 | Uncharacterised DC. Bone marrow failure and nail dystrophy. Sibling of *DKC*28 | None |
| DC 30 | F | 43 | *TERT* variant Ile 1130 Val. Dry skin. | None |
| DC 31* | F | 34 | *TERC* Mutation. Bone marrow failure | None |
| DC32 | F | 36 | *TERC* Mutation. Bone marrow failure and skin abnormalities. | Danazol 100mg daily |
| DC 33 | M | 69 | Uncharacterized DC. Leukopenia and nail dystrophy | None |
| DC 34 | M |  | *TERT* variant (Arg865His). Past history of Hodgkin’s lymphoma | None |

F = Female M = Male * Father son and daughter + Father and son ++ Mother and son ** Aunt and nephew

All *TERC*, *TERT,* and *RTEL* mutations are heterozygous and all *DKC1* mutations are hemizygous (X-linked). *NOP10* mutations are biallelic*.*

**Table 2. Age and Gender of control subjects.**

| Subject Number | Gender | Age |
| --- | --- | --- |
| C1 | F | 33 |
| C2 | M | 63 |
| C3 | F | 27 |
| C4 | M | 61 |
| C5 | M | 56 |
| C6 | M | 35 |
| C7 | F | 28 |
| C8 | M | 49 |
| C9 | M | 26 |
| C10 | M | 31 |
| C11 | M | 23 |
| C12 | M | 29 |
| C13 | M | 64 |
| C14 | F | 92 |
| C15 | F | 77 |
| C20 | F | 26 |
| C21 | F | 27 |
| C22 | F | 28 |
| C23 | M | 35 |
| C24 | M | 34 |
| C25 | F | 26 |
| C26 | M | 35 |
| C27 | F | 35 |
| C28* | M | 23 |
| C29 | F | 29 |
| C30* | F | 38 |
| C31 | F | 27 |
| C32 | F | 38 |
| C33 | M | 28 |
| C34 | M | 27 |
| C35 | Unknown | 54 |
| C36 | Unknown | 20 |
| C37 | Unknown | 38 |

**Table 3A. Rank Test P values of DC Patients (n=29) versus normal donors (n =30.)**

| P Value | Citrate | Aconitate | Isocitrate | Malate | Succinate | Lactate | Pyruvate |
| --- | --- | --- | --- | --- | --- | --- | --- |
| All n=29 | 0.08 | **0.03** | **0.0007** | **0.0005** | **0.0008** | **0.0003** | **0.0000007** |
| Severe bone marrow Symptoms  n =12 | 0.19 | 0.21 | **0.04** | **0.002** | **0.001** | **0.006** | **0.00002** |
| Asymptomatic  or mild bone marrow symptoms  n =17 | 0.15 | 0.09 | **0.001** | **0.007** | **0.03** | **0.002** | **0.0001** |
| Mucocutaneous  Symptoms (n =15) | 0.14 | **0.03** | **0.008** | **0.002** | **0.0006** | **0.008** | **0.000004** |
| Males only** | 0.20 | 0.06 | **0.01** | **0.006** | **0.01** | **0.008** | **0.0001** |
| *TERC* Mutants only*** | 0.40 | 0.56 | **0.005** | **0.006** | **0.02** | **0.01** | **0.0002** |
| *TERT* Mutants Only**** | 0.14 | **0.048** | **0.003** | **0.008** | **0.01** | **0.02** | **0.003** |
| *DKC1* Mutants Only***** | 0.33 | ND | 0.5 | 0.16 | **0.04** | 0.06 | **0.002** |

**Rank Test P values of DC Patients (n=27) versus normal donors (n =19)**

|  | Glucose |
| --- | --- |
| All (n=27) | 0.25 |

|  | 7-MG | C-MT | Urate | IL-6* | OAA | IL-1α | Alpha  KG | Lactate:Pyruvate Ratio |
| --- | --- | --- | --- | --- | --- | --- | --- | --- |
| All n =29 | 0.50 | 0.23 | 0.67 | 0.11* | 0.06 | 0.06+ | 0.39 | **0.005** |
| Severe bone marrow Symptoms n = 12 | 0.96 | 0.30 | 0.44 | ND | 0.09 | ND | 0.52 | **0.004** |
| Asymptomatic  or mild bone marrow symptoms  n =17 | 0.34 | 0.35 | 1.0 | ND | 0.16 | ND | 0.45 | 0.08 |
| Mucocutaneous  Symptoms (n =15) | 0.50 | 0.72 | 0.94 | ND | 0.12 | ND | 0.36 | **0.002** |
| Males only** | **0.05** | 0.22 | 0.15 | ND | 0.26 | ND | 0.83 | **0.004** |
| *TERC* Mutants Only*** | 0.56 | **0.03** | 0.44 | ND | 0.18 | ND | **0.03** | **0.02** |
| *TERT* Mutants Only**** | 0.97 | 0.59 | 0.91 | ND | 0.23 | ND | 0.20 | 0.33 |
| *DKC1* Mutants Only***** | 0.22 | 0.77 | 0.24 | ND | 0.27 | ND | 0.42 | 0.12 |

*15 DC Patients versus 20 controls ** 20 DC patients versus 14 controls. *** 12 DC patients with *TERC* mutations versus 30 control subjects. **** 7 DC patients with *TERT* variants/mutations versus 30 control subjects. ***** 6 DC patients with *TERT* variants/mutations versus 30 control subjects. + 15 DC patients and 10 severely symptomatic so far. Only one patient with reliably detectable levels. Values in bold indicate statistically significant < 0.05 by the Wilcoxon-Mann-Whitney Test. Green = reduced compared to normal; red = elevated compared to normal.

**Table 3B. Rank Test P values of DC Patients (n=29) versus normal donors (n =30) normalised to control average and corrected for false discovery rate.**

| P Value | Citrate | Aconitate | Isocitrate | Malate | Succinate | Lactate | Pyruvate |
| --- | --- | --- | --- | --- | --- | --- | --- |
| All n=29 | 0.13 | **0.03** | **0.001** | **0.003** | **0.003** | **0.001** | **0.0000006** |

**Table 4. Linear regression P values of plasma levels of metabolites versus LAATL in DC patients.**

| P Value | Citrate | Aconitate | Isocitrate | Malate | Succinate | Alpha KG |
| --- | --- | --- | --- | --- | --- | --- |
| All n=19 | **0.01** | 0.92 | 0.26 | **0.03** | 0.16 | 0.44 |
| Severe Symptoms n =11 | 0.14 | 0.98 | 0.31 | 0.18 | 0.99 | 0.42 |
| *TERC* mutations only  n =12 | **0.05** | 0.61 | 0.50 | **0.03** | 0.78 | 0.52 |

|  | Lactate | Pyruvate | 7-MG | C-MT | Urate | IL-6* | OAA |
| --- | --- | --- | --- | --- | --- | --- | --- |
| All n =19  * n = 9 | 0.22 | 0.17 | 0.99 | 0.98 | 0.66 | 0.28 | 0.25 |
| Severe Symptoms n =11 | 0.32 | 0.45 | 0.82 | 0.88 | 0.95 | ND | 0.97 |
| *TERC* mutations only  n =12 | 0.26 | 0.29 | 0.66 | 0.58 | 0.85 | ND | 0.52 |

**Table 5. Relationship of Citrate to other metabolites by linear regression analysis.**

|  | **Isocitrate** | **Aconitate** | **Malate** | **Succinate** |
| --- | --- | --- | --- | --- |
| **Normal Subjects** | **0.002** | **0.0007** | 0.11 | 0.57 |
| **DC Patients** | **5.83 x 10^-6^** | **6.63 x 10^-5^** | **9.1 x 10^-5^** | **0.004** |

|  | **Oxaloacetic Acid** | **Alpha Ketoglutarate** | **Lactate** | **Pyruvate** |
| --- | --- | --- | --- | --- |
| **Normal Subjects** | 0.41 | 0.28 | 0.16 | 0.13 |
| **DC Patients** | 0.52 | 0.15 | 0.08 | 0.51 |

|  | **Lactate:Pyruvate Ratio** | **C-mannosyl tryptophan** | **7-methyl guanine** | **Urate** |
| --- | --- | --- | --- | --- |
| **Normal Subjects** | 0.59 | **0.0002** | **0.002** | 0.13 |
| **DC Patients** | 0.82 | 0.11 | **0.02** | 0.88 |

|  | **Glucose** |
| --- | --- |
| **Normal Subjects (n =19)** | 0.83 |
| **DC Patients (n =21)** | 0.44 |

**Table 6. Linear regression P values of plasma levels of metabolites versus plasma levels of IL-6 in *DC* patients (n = 15 by GC/MS and LC/MS).**

| P Value | Citrate  GC/MS | Citrate  LC/MS | Aconitate | Isocitrate | Malate | Succinate |
| --- | --- | --- | --- | --- | --- | --- |
| R^2^ | 0.30 | 0.4046 | 0.31 | 0.5675 | 0.38 | 0.28 |
| P | **0.01** | 0.06 | 0.10 | **0.02** | 0.08 | 0.42 |

|  | 7-MG | C-MT | Urate | Alpha KG | OAA | Lactate | Pyruvate | Lactate:Pyruvate  Ratio |
| --- | --- | --- | --- | --- | --- | --- | --- | --- |
| R^2^ | 0.7873 | 0.1676 | 0.025 | 0.0007 | 0.039 | 0.09 | 0.018 | 0.007 |
| P | **0.008** | 0.21 | 0.65 | 0.97 | 0.54 | 0.28 | 0.70 | 0.76 |

**Table 7. Linear regression P values of plasma levels of metabolites versus DC clinical indicators of aplastic anaemia in DC patients.**

**Metabolites versus White Blood Cell Count.**

| P Value | Citrate | Aconitate* | Isocitrate | Malate | Succinate | Alpha Ketoglutarate |
| --- | --- | --- | --- | --- | --- | --- |
| All n=27 | 0.96 | 0.28 | 0.49 | 0.45 | 0.79 | 0.20 |

|  | 7-MG | C-MT | Urate | OAA | Lactate | Pyruvate | Lactate:Pyruvate Ratio |
| --- | --- | --- | --- | --- | --- | --- | --- |
| All n =27 | 0.61 | 0.96 | 0.10 | 0.87 | 0.83 | 0.58 | 0.43 |

**Metabolites versus Platelet Count.**

| P Value | Citrate | Aconitate* | Isocitrate | Malate | Succinate | Alpha Ketoglutarate |
| --- | --- | --- | --- | --- | --- | --- |
| All n=27 | 0.72 | 0.38 | 0.46 | 0.75 | 0.56 | 0.64 |

|  | 7-MG | C-MT | Urate | OAA | Lactate | Pyruvate | Lactate:Pyruvate Ratio |
| --- | --- | --- | --- | --- | --- | --- | --- |
| All n =27 | 0.67 | 0.67 | 0.22 | 0.72 | 0.92 | 0.68 | 0.66 |

**Metabolites versus Haemoglobin.**

| P Value | Citrate | Aconitate* | Isocitrate | Malate | Succinate | Alpha Ketoglutarate |
| --- | --- | --- | --- | --- | --- | --- |
| All n=27 | 0.55 | 0.25 | 0.24 | 0.53 | 0.83 | 0.68 |

|  | 7-MG | C-MT | Urate | OAA | Lactate | Pyruvate | Lactate:Pyruvate Ratio |
| --- | --- | --- | --- | --- | --- | --- | --- |
| All n =27 | 0.67 | 0.93 | 0.93 | 0.88 | 0.19 | 0.75 | 0.10 |

**Metabolites versus Mean Corpuscular Volume.**

| P Value | Citrate | Aconitate* | Isocitrate | Malate | Succinate | Alpha Ketoglutarate |
| --- | --- | --- | --- | --- | --- | --- |
| All n=27 | 0.39 | 0.46 | 0.57 | 0.34 | 0.70 | 0.83 |

|  | 7-MG | C-MT | Urate | OAA | Lactate | Pyruvate | Lactate:Pyruvate Ratio |
| --- | --- | --- | --- | --- | --- | --- | --- |
| All n =27 | 0.22 | 0.81 | 0.16 | 0.54 | 0.79 | 0.30 | 0.34 |

* n = 26

**Table 8. The effect of age and gender on LC/MS metabolites.**

A. **Normal Subjects.**

| P Value | Citrate | Aconitate | Isocitrate | Malate | Succinate | Lactate | Pyruvate |
| --- | --- | --- | --- | --- | --- | --- | --- |
| Age n= 30 | 0.37 | 0.89 | 0.72 | 0.52 | 0.43 | 0.11 | 0.54 |
| Gender  n =27  (14 male 13 female) | 0.86 | 0.81 | 0.83 | 0.93 | 0.84 | 0.37 | 0.11 |

|  | 7-MG | C-MT | Urate | OAA | Alpha  KG | Lactate:Pyruvate Ratio |
| --- | --- | --- | --- | --- | --- | --- |
| Age n = 30 | 0.14 | 0.79 | 0.11 | 0.19 | 0.34 | 0.92 |
| Gender n =27 (14 male 13 female) | 0.43 | 0.98 | 0.22 | 0.21 | 0.10 | **0.02** |

**B. DC Patients.**

| P Value | Citrate | Aconitate* | Isocitrate | Malate | Succinate | Lactate | Pyruvate |
| --- | --- | --- | --- | --- | --- | --- | --- |
| Age n= 28 | 0.22 | 0.37 | 0.96 | **0.03** | 0.92 | 0.54 | 0.21 |
| Gender  n =29(*28)  (19; *18 male 10 female) | 0.47 | 0.47* | 0.20 | 0.12 | 0.38 | **0.02** | 0.85 |
| Danazol (n = 6; * n = 5) | 0.80 | 0.06* | 0.15 | 0.09 | **0.0008** | **0.02** | **0.0006** |
| No Danazol (n = 23) | **0.05** | 0.07 | **0.0007** | **0.0006** | **0.01** | **0.001** | **0.000009** |
| Danazol v No Danazol | 0.67 | 0.27 | 0.83 | 0.48 | **0.01** | 0.75 | 0.75 |

|  | 7-MG | C-MT | Urate | OAA | Alpha  KG | Lactate:Pyruvate Ratio |
| --- | --- | --- | --- | --- | --- | --- |
| Age n = 28 | 0.50 | 0.30 | 0.94 | 0.82 | 0.48 | 0.92 |
| Gender n =29  (19 male 10 female) | **0.05** | 0.22 | 0.46 | 0.20 | 0.89 | 0.22 |
| Danazol (n = 6) | 0.50 | 0.52 | 0.44 | 0.14 | 0.87 | 0.25 |
| No Danazol (n = 23) | 0.60 | 0.25 | 0.86 | 0.11 | 0.29 | **0.005** |
| Danazol v No Danazol | 0.67 | 1.0 | 0.71 | 0.59 | 0.33 | 0.33 |

**Tale 9. Effect of sample time and batch on metabolites in normal subjects.**

| P Value | Citrate | Aconitate | Isocitrate | Malate | Succinate | Lactate | Pyruvate |
| --- | --- | --- | --- | --- | --- | --- | --- |
| Batch 1 (p.m. n =15) v  Batch 2 (a.m. n =15) | 0.6 | 0.85 | 0.44 | 0.66 | 0.95 | 0.12 | 0.27 |

|  | 7-MG | C-MT | Urate | OAA | Alpha  KG | Lactate:Pyruvate Ratio |
| --- | --- | --- | --- | --- | --- | --- |
| Batch 1 (p.m. n =15) v  Batch 2 (a.m. n =15) | 0.52 | 0.42 | 0.10 | 0.85 | 0.37 | 0.11 |

**Table 10. Average variation between repeat samples of 8 DC patients.**

| Metabolite | Average (%) Standard Deviation  between repeat DC samples ( n =16) |
| --- | --- |
| Oxaloacetic Acid | 28.2 |
| 7-methylguanine | 19.1 |
| C-mannosyl tryptophan | 5.7 |
| Urate | 32.0 |
| Citrate | 10.5 |
| Isocitrate | 15.4 |
| Aconitate | 13.5 |
| Malate | 21.4 |
| Succinate | 18.4 |
| Lactate | 22.6 |
| Pyruvate | 18.5 |
| Lactate:Pyruvate Ratio | 1.0 |

**Table 11. The effect of low dose Danazol treatment in DC patients (n =6) v no treatment (n =21).**

|  | Citrate | Aconitate | Isocitrate | Malate | Succinate | Lactate | Pyruvate |
| --- | --- | --- | --- | --- | --- | --- | --- |
| P | 0.67 | 0.29 | 0.83 | 0.48 | **0.01** | 0.75 | 0.75 |

|  | 7-MG | C-MT | Urate | OAA | Alpha  KG | LPR |
| --- | --- | --- | --- | --- | --- | --- |
| P | 0.67 | 1.0 | 0.71 | 0.59 | 0.33 | 0.52 |
